# Pigeons rapidly transmit information about predator location through gaze-following

**DOI:** 10.1101/2025.10.29.685277

**Authors:** Mathilde Delacoux, Michael Chimento, Fumihiro Kano

**Affiliations:** Centre for the Advanced Study of Collective Behaviour, University of Konstanz; Universitätsstraße 10, 78464 Konstanz, Germany; Department of Biology, University of Konstanz; Universitätsstraße 10, 78464 Konstanz, Germany; Department of Collective Behavior, Max Planck Institute of Animal Behavior; Am Obstberg 1, 78315 Radolfzell, Germany; International Max Planck Research School for Quantitative Behavior, Ecology and Evolution; Am Obstberg 1, 78315 Radolfzell, Germany; Department of Evolutionary Biology and Environmental Studies, University of Zurich, Zurich, Switzerland; Institute for Advanced Study, Kyushu University; Fukuoka, Japan

**Keywords:** Anti-predatory vigilance, Animal cognition, Collective detection, Social transmission, Visual attention, Network-based diffusion analysis, 3D posture tracking, Columba livia domestica

## Abstract

Animals in groups benefit from the social transmission of key information, such as when a predator appears. However, ambiguous social cues may lead to false alarms. Gaze-following—aligning gaze direction with others— could mitigate this risk by allowing individuals to verify predator location prior to costly escape. However, its role in collective vigilance remains unclear. Here, we experimentally show that pigeons (*Columba livia domestica*) foraging in flocks use gaze-following to detect predator location, before initiating evasion. We used high-resolution head tracking to reconstruct visual fields and estimate foveal orientation toward the predator stimulus. We quantified social transmission of predator location using a dynamic multi-network Bayesian diffusion model. In addition to personal information, pigeons responded to the duration of conspecific gaze toward the predator, rather than alternative cues, such as locomotion or head-up vigilance. Gaze following can act as an early mechanism for transmitting predator-related information in a flock.

## Introduction

Social information transmission is a key advantage of group living [1–4], especially in uncertain or high-stakes contexts [5–9]. During collective predator vigilance, individuals often initiate evasive action based on social cues, such as others’ escape, before detecting threats [10–14]. Yet without explicit signals [15, 16], such cues can be ambiguous and prone to costly false alarms [17–20], diminishing the benefits of collective vigilance [8]. Gaze following (aligning one’s gaze with that of another), a socio-cognitive skill widespread across species [21–30], may reduce this ambiguity by allowing individuals to localize threats and avoid unnecessary escapes. Although this function has long been hypothesized, empirical support has remained limited.

During collective vigilance, birds frequently interrupt ongoing activities such as foraging with brief bouts of threat assessment[31–34]. Yet the processes through which individuals detect and transmit information during these moments remain poorly understood, partly because addressing this question requires fine-grained measurements of the social and environmental cues available within their visual fields[35]. We asked whether gaze following, or aligning one’s gaze with that of another, functions to transmit specific information about predator location during collective foraging. Gaze following has been studied extensively in terms of its cognitive mechanisms [25, 27, 36–39], phylogeny [21–30], and development [27, 29, 30, 40–44], and the idea that it may serve to share information during vigilance has long been proposed yet has never been empirically demonstrated.

Using a recently developed fine-scale behavioural tracking system for pigeons based on motion capture, visual fields, including projections of the foveae (the retinal regions of high-acuity vision) were reconstructed from head orientation [45, 46]. Using the same tracking system, pigeons have also been shown to follow conspecific head orientation toward specific targets in controlled gaze-following paradigms based on conventional avian gaze-following designs [25, 27, 28]. In addition, evasive flight responses in pigeons are socially contagious [32, 47], making them well suited for investigating social transmission of predator-related information.

To model information transmission, we fit dynamic, multi-network Bayesian transmission models (R package STbayes) [48], which integrate recent advances in network-based diffusion analysis (NBDA) [49], multilayer network modelling [50], and visual sensory network analysis of collective information flow [20, 51, 52]. This approach allowed us to account for the dynamically changing structure of networks and the multiple pathways through which information might propagate.

Our rationale was that if gaze following occurs during predator vigilance, a pigeon that orients its gaze toward the predator (henceforth an “informed individual”) can inadvertently provide a cue to flock-mates indicating the direction of the threat. In order to avoid false positives, we define a threshold of gazing longer than *>* 300ms at the predator for a bird to become informed of its location. An observation of this cue by an “uninformed individual” would then increase the probability that the uninformed individual would subsequently also gaze toward the predator (Fig 1a). Thus, we predicted that the probability of an uninformed individual foveating at the predator at time *t* should be best fit by a model that specifically includes information about the magnitude of network connections to already informed individuals (i.e. emitting gaze cues, Fig 1f).

**Figure 1.**
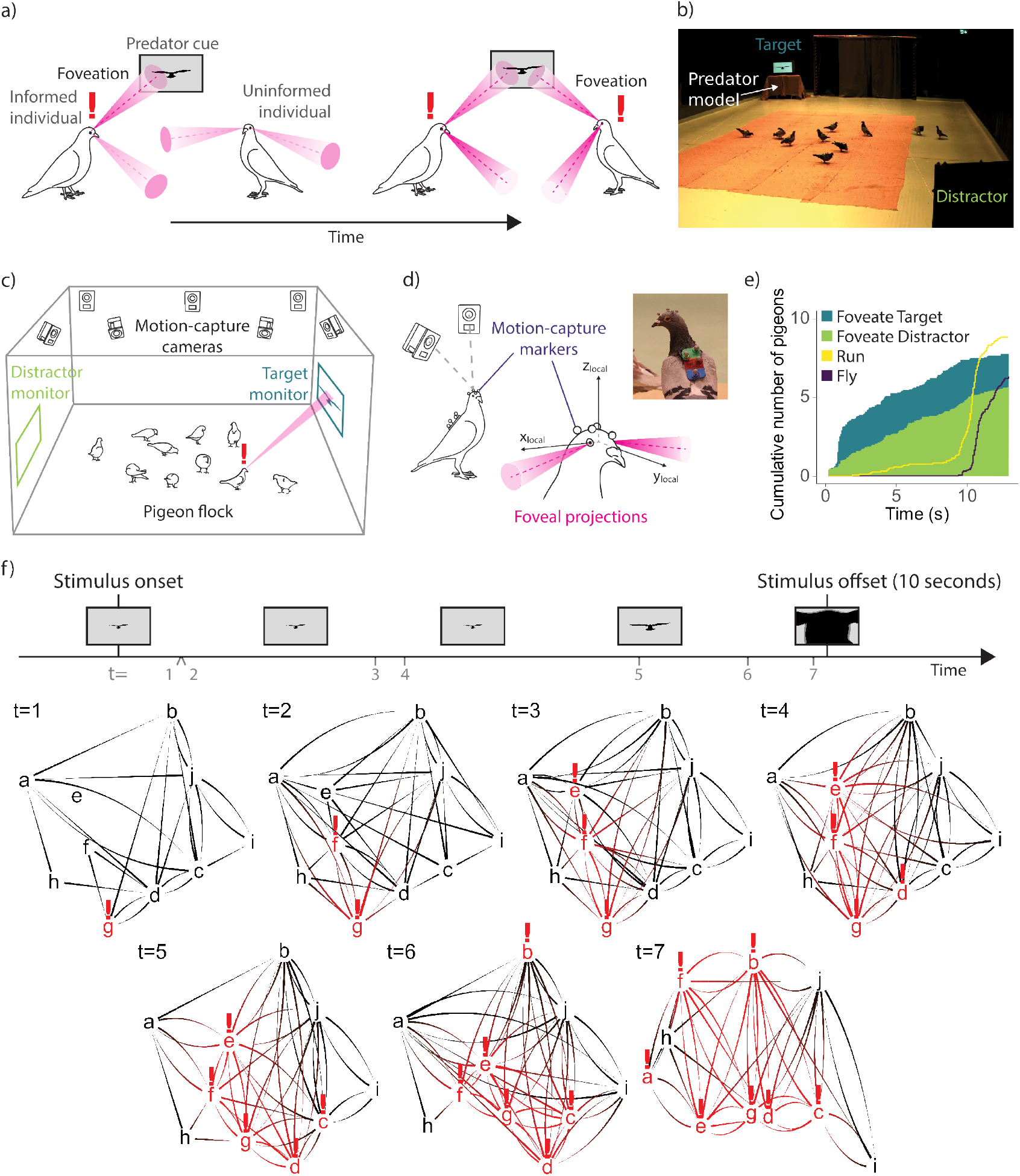
Experimental system demonstrating social transmission of predator information via gaze following. a) Definition of “being informed”. A pigeon was defined as “foveating” when one of its gaze vectors intersected with a monitor or a conspecific. An “uninformed” individual was considered “informed” after its first sustained foveation on the target monitor. b) The experimental room with a flock of foraging pigeons. The “target” monitor displayed the looming predator stimulus, while the “distractor” monitor remained blank. c) Schematic representation of the experimental room and motion-capture system (not at scale). d) Reconstruction of “gaze” in a pigeon. Each pigeon wore motion-capture markers on its head and back. Foveal projections were reconstructed as cones within a head-centered coordinate system. e) Cumulative number of individuals that foveated on the target and distractor monitors, and that made an evasion response (running away or flying) over the course of the simulated predation event. f) Illustrative example of social transmission of predator information via gaze following across a dynamic visual field network. We divided each trial into discrete periods based on the sequence of “first foveations”, with each period ending when one new pigeon “detected” the simulated predator. At each timestep, informed individuals are shown in red with an exclamation mark. Edges connections between individuals are calculated from continuous 3D-tracking data of head orientation and body position. The direction of information flow is indicated by the edge shape: the broader end points to the transmitting individual, the narrower end to the receiver. The data underlying this Figure can be found in https://doi.org/10.17617/3.AWIUN8.

## 1 Results

We subjected flocks of 10 pigeons (from a total of 20 pigeons) to 24 simulated predator events, for a total of 240 observations. The flocks were arranged such that the composition was different across all 12 trials and each pigeon was paired with all other individuals at least one time. All pigeons were naïve to predator presentations prior to the experiment and experienced a total of 12 predator presentations. In each trial, pigeons naturally foraged on scattered grain until a looming predator stimulus was presented. A motion-capture system tracked fine-scale head and body movements within a large space (Fig 1b-c). We reconstructed individuals’ visual fields and foveal projections (Fig 1d). Gaze was operationalized as a conservative 10° foveal projection cone derived from anatomical calibration, accommodating the relatively limited range of eye movements in freely moving pigeons (typically *<*5°) [53]. Validation work confirmed that the modelled foveal projection captures the spatial extent of high-acuity vision, as pigeons reliably orient their lateral foveae (not binocular field) toward behaviorally relevant targets, including objects, food, conspecifics, and predator stimuli [46, 47].

Each simulated predator event consisted of a looming raptor shadow displayed on a monitor (Fig 1b-c), followed by a plastic raptor model emerging from a hidden compartment located directly beneath the monitor and moving across the room on a pulley system. Two monitors were positioned on opposite sides of the experimental room: one displayed the looming predator stimulus (“target” monitor), while the other (“distractor” monitor, Fig 1b-c) did not, instead showing a light grey screen. The locations of the target and distractor monitors were counterbalanced across events. The majority of individuals in the flock foveated on the target monitor (median 8 birds) and, to a lesser extent, on the distractor monitor (median 6 birds) by the end of each predator event (Fig 1e). Although some habituation was observed regarding the flight response to the plastic model predator, pigeons consistently looked at the preceding predator looming cue across trials (S1 Fig), and we found no effect of trial in a posthoc test (see Materials and Methods).

With this data, we aimed to answer three questions: 1) Did pigeons use social information to detect the predator? 2) If so, what were the pathways of social transmission, and 3) what cues were important for transmission? To answer these questions, we designed a set of hierarchical, Bayesian network-based diffusion analysis (NBDA) models representing alternative hypotheses about each question [48]. NBDA models are widely used in the animal culture literature to determine whether social transmission of novel behaviors is occurring, how strong its effects are, and the exact pathways of information transfer [49, 54, 55]. These models predicted the times that each individual became informed of the predators’ location, defined as the first time they foveated on the predator. Predictors included their dynamic network connections to other individuals, and those individuals’ dynamic cue states as transmission weights (e.g. foveating on predator or not). The model then estimated whether connections to informed individuals increase the rate at which others become informed. Fig 2 illustrates the general structure of our main multi-network gaze-following model, which estimates a rate of becoming informed as a function of an asocial baseline detection rate and a social transmission rate. A strength of social transmission was estimated for each network of interest, and is interpreted as the degree to which connections to informed, cueing (e.g. foveating at predator stimulus) flock-mates increases the probability of becoming informed relative to the asocial rate. We also included whether individuals were in a “head down” position as a variable that could affect asocial detection and social transmission, as pigeons may be less attentive to their environment when actively foraging. We performed model comparison to test different alternative hypotheses using LOO-PSIS, a measure of out-of-sample predictive fit [56]. By comparing models with different likelihood formulations (e.g. with and without social transmission), different network and different cue assumptions, we could infer the answers to the questions above.

**Figure 2.**
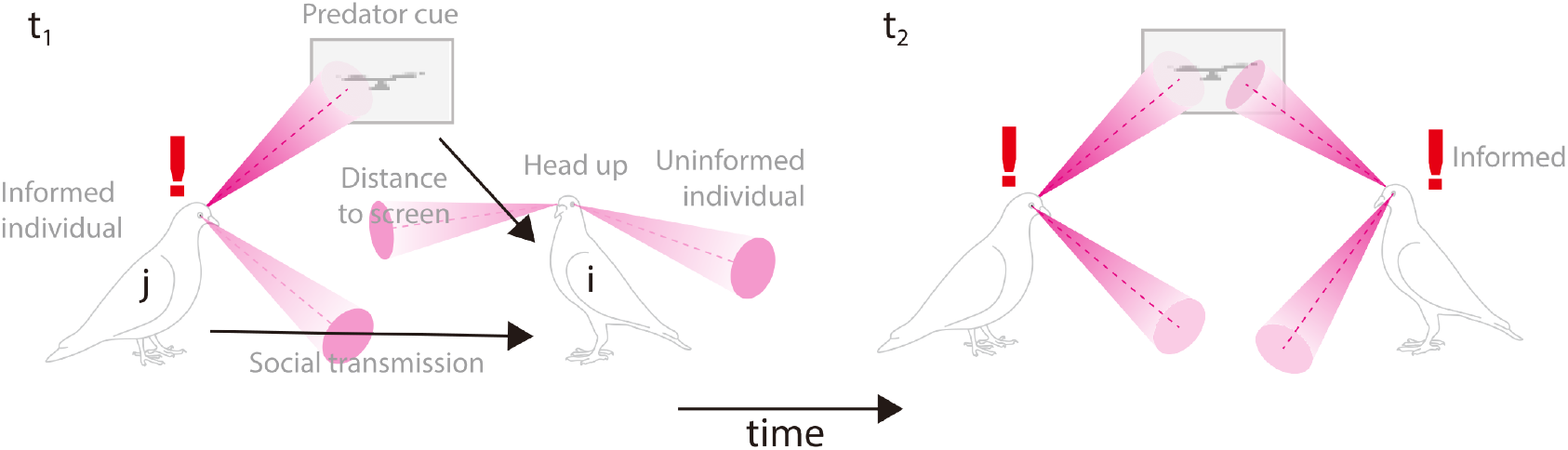
Model of gaze following. We model the probability of individual *i* becoming informed as a function of a baseline asocial rate of detection, a social transmission rate, *i*’s proximity to the monitor, and the proportion of time *i* spent in a “head up” posture. Social transmission was defined as the sum of network connections to informed conspecifics (e.g. *j*) in each timestep, multiplied by whether or not *j* was emitting a cue at that timestep. Thus, non-cueing individuals did not provide information to uninformed individuals. In the gaze following model, the cue was defined as foveating on the target.

### Did pigeons use social information to detect the predator?

To address whether social transmission plays an important role in predator detection, we began by fitting a set of asocial models that assumed no social transmission. Based on previous findings [52], we expected pigeons positioned closer to the simulated predator or with a clearer view of it to become informed more quickly. We also expected pigeons in a “head-down” position (likely foraging for grains and not attending to their environment) to become informed less quickly. To test this, we compared models with and without these effects on the baseline detection rate. The model including effects of visual field coverage by the monitor and whether pigeons were head down was most predictive (S1 Table; similar results were observed for the distractor monitor S2 Table). The larger the target monitor appeared within a pigeon’s visual field, the more likely the bird was to become informed (*β*_*M*_ (95% HPDI) = 2.889(2.159, 3.626)). Additionally, a head-down state decreased the likelihood of becoming informed (*β*_*HD*_ = *−*2.474( *−*2.993, *−*1.981)). Accordingly, we included these variables in all subsequent models. Next, we compared the asocial model to our main multi-network social (gaze following) model. This model estimated the relative strength of social transmission between individuals across 4 different network types: visual field, voronoi, inverse distance and foveation (Fig 3a, S3 Table). The model was significantly more predictive than the best-fitting asocial model (Δ*LOOIC* = *−*31.17 ( *−*44.09, *−*18.25), **control analysis A**, Fig 4), indicating that social transmission played a crucial role in the detection of the predator’s location. Although a majority of detections occurred asocially, the model estimated that 28.1% detections (S4 Table) were triggered by social information transmission through the visual field in the multi-network model. In a single-network model considering visual field as only means of transmission, 35.8% of events occurred due to social transmission (see below and S6 Table).

**Figure 3.**
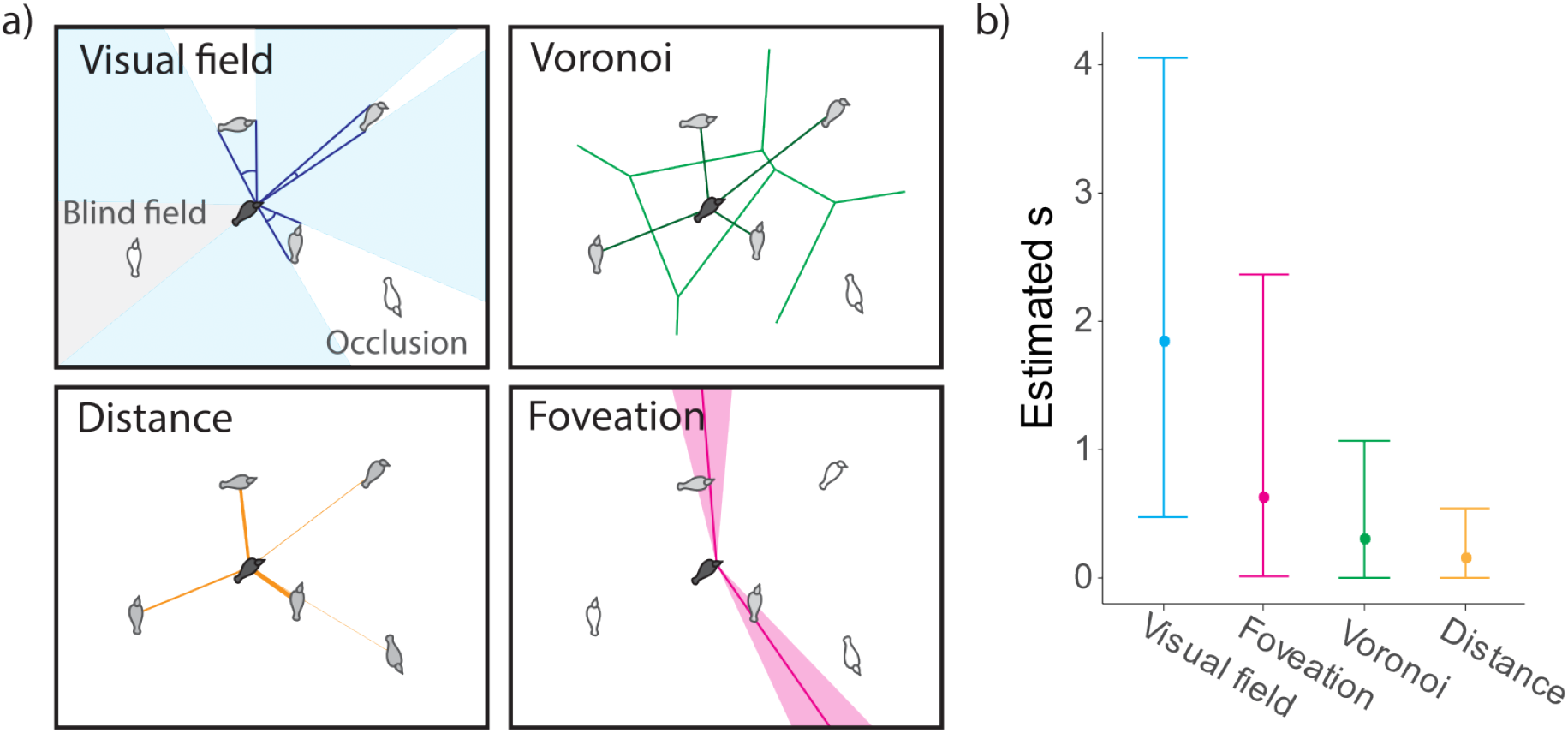
Networks used in the transmission analysis. a) Visual representation of the 4 different types of networks used for the multi-network model. b) Estimated *s* of the different networks with their 95% CI. The data underlying this Figure can be found in https://doi.org/10.17617/3.AWIUN8.

**Figure 4.**
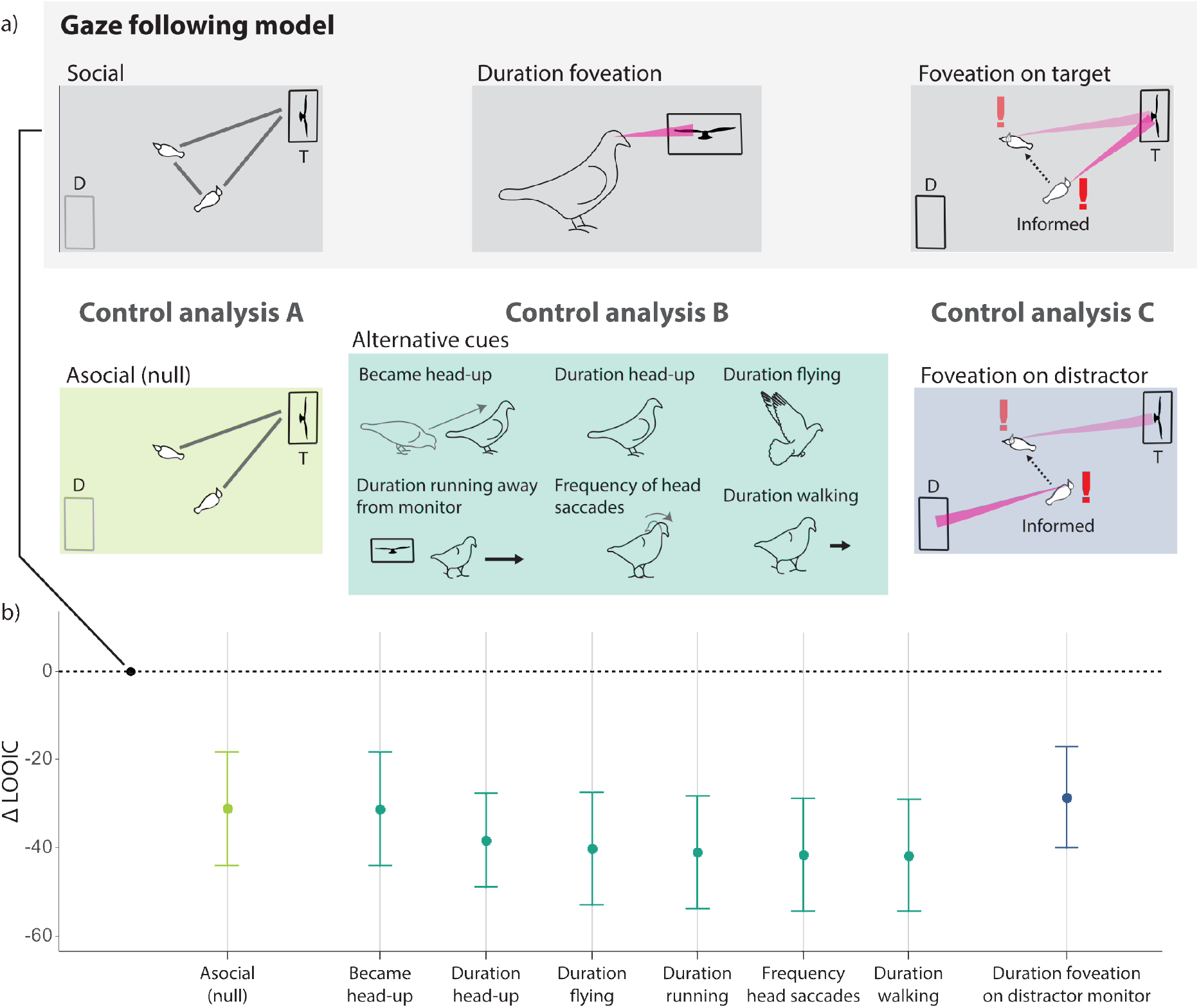
Control analysis results. a) Schematic overview of the control analyses. Control Analysis A compares the Gaze following (main) model, which includes social transmission, with an asocial (null) model that does not include it. Control Analysis B compares the Gaze following model, which uses the duration of foveation on the target monitor, with models using alternative behavioural cues (e.g., head-up posture, locomotion). Control Analysis C compares models using foveation on the target monitor (Gaze following) versus foveation on the distractor monitor as the social cue. T = Target monitor; D = Distractor monitor. b) Model comparison results. Each dot represents the Δ*LOOIC* (Leave-One-Out Information Criterion) score of a model, with the error bars indicating standard error of the difference. A negative Δ*LOOIC* indicates worse model fit as evaluated by out-of-sample predictiveness. The horizontal dotted line marks the Δ*LOOIC* of the Gaze following model; models with error bars entirely below this line indicate models that were significantly less predictive than the Gaze following model. The data underlying this Figure can be found in https://doi.org/10.17617/3.AWIUN8.

### What was the pathway of social transmission?

In the multi-network model, the visual field network received the highest estimated strength of social transmission 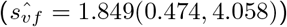, while all other networks received comparatively low estimates whose credible intervals approached 0 (Fig 3b, S4 Table). The quantity *s* describes the relationship between social and asocial rates, which both additively contribute to the overall detection rate. Thus, the model estimate can be interpreted here as meaning that for each cueing bird in the visual field, there is a 95% chance that the overall rate of detection is approximately between 1.5 *−* 5 times faster than if no informed birds were in the visual field.

We then compared the multi-network model with single-network models, including each one of the four tested networks. The single-network visual field model and the multi-network model were the most predictive, with no detectable statistical difference as measured by Δ*LOOIC* (S5 Table, results of the best-fitting single network model in S6 Table). The superiority of the visual field network over the foveation and proximity (inverse distance and Voronoi) networks suggests that pigeons primarily detected conspecific gaze cues through peripheral vision.

### What cues were responsible for social transmission?

We compared the main gaze-following model to alternative models that assumed different cues defining the dynamic transmission weights *w*_*jt*_ (Fig 4a). We used these comparisons to infer whether pigeons responded to other vigilance-related behaviours (**control analysis B**). The head-up posture and scanning behaviour (common avian vigilance indicators [35]) were measured by the duration spent in a head-up posture, the number of head-down to head-up transitions, and the frequency of head-scan saccades. We also assessed responses to locomotion cues, including walking, running away from the monitor, and flying. None of the models based on these alternative cues were more predictive than the main gaze-following model (S5 Table, Fig 4b).

We then asked whether pigeons were responding to the act of conspecific “staring”, irrespective of gaze direction, and only subsequently detecting the predator as a salient stimulus. To test this, we constructed an alternative model (**control analysis C**) that predicted foveation on the target monitor using conspecific gaze directed toward the distractor monitor as a cue. If pigeons were simply reacting to gaze behaviour irrespective of direction, this model should perform comparably to the main model. Instead, it performed significantly worse (Fig 4b), indicating that pigeons responded specifically to gaze directed at the target.

Finally, we considered whether gaze following was triggered by the combination of conspecific gaze and a salient predator stimulus, rather than by gaze cues alone. To test this, we constructed models that predicted foveation on the distractor monitor, using conspecific gaze directed either at the distractor (which did not display the predator stimulus) or at the target monitor as cues (**control analysis D**). This analysis, conceptually similar to control analysis C, tested whether pigeons also followed the gaze of conspecifics toward the distractor monitor (a non-threatening location). The model using conspecific gaze directed at the distractor provided a better fit than the model using gaze directed at the target when predicting gaze toward the distractor (ΔLOOIC = *−*24.05 [ *−*41.44, *−*6.67]; S7 Table, S2 Fig; results of the best-fitting multi-network and single network models in S8, S9 Tables). These results indicate that pigeons can use conspecific gaze cues alone, even when no predator is present at that location.

## Discussion

We show that gaze following serves as an early pathway for transmitting predator location information within a flock without active signaling involved. It was specifically triggered by conspecifics’ gaze toward the predator, not by their locomotion, head-up vigilance, or gaze directed elsewhere. Although it cannot be certain that pigeons interpreted the looming cue on the monitor as a predator, this indicates that spatial information about a potential threat can be transmitted through this mean. Pigeons relied primarily on their peripheral visual fields rather than foveal vision to respond to flock-mates’ gaze cues, suggesting that gaze following does not require close visual attention to conspecifics (i.e., even during foraging). Notably, individuals occasionally followed gaze directed toward a non-threatening location (the distractor monitor), responding solely to conspecific gaze cues. Yet despite these deviations, by the end of each predator presentation more pigeons had foveated on the target than on the distractor (Fig 1e), indicating that the flock collectively distinguished between the two.

During collective vigilance, the social cues available to individuals are often ambiguous, and responding to false alarms can result in missed opportunities for other activities (e.g., foraging, courting) or in costly, unnecessary escape responses [8, 18]. To limit the spread of false alarms, individuals may require a threshold number of escaping conspecifics before initiating their own escape [17, 19, 20, 33, 52]. Keeping distance from others and reducing group cohesion can further buffer against the propagation of weak or isolated alarms [20]. Individuals may also increase their vigilance in response to the alertness or escape behaviour of others [31, 33, 34, 57, 58], or dynamically adjust their sensitivity to alarms according to their perceived reliability [59, 60]. Balancing social information use with personal assessment therefore remains a central challenge for individuals engaged in collective vigilance.

Our findings highlight how individual attention and group coordination mitigate the collective challenges of predator detection. Although the majority of detections occurred through individual detections, a substantial proportion (approximately 1/3) depended on the transmission of social information. Following conspecific gaze likely provides three key benefits: (1) it increases the chance of detecting a predator earlier than would be possible individually; (2) it enables direct assessment of the threat after foveation by providing information about its location; (3) it requires minimal movement and time investment; and (4) it only relies on inadvertent cues given by conspecifics and not on active signaling. Together, this strategy enhances detection while reducing the costs of missed opportunities, unnecessary energy expenditure, and false alarms triggered by social cues.

Gaze following has been extensively studied across species [22, 25, 37, 61]. It emerges early in development [40–44] and is widely documented in mammals [21–24], birds [25–28], and likely also in reptiles [27, 29] and fishes [30]. While gaze following can involve slower, perspective-taking processes [25, 27, 36], its simplest form is rapid and reflexive [37–39]. This reflexive mode may enable gaze following to function effectively in high-stakes contexts such as predator detection, where both speed and accuracy are critical. While we modelled social transmission as a linear function of exposure to other informed individuals who were emitting cues, it is possible that social transmission follows a non-linear rule, such as a frequency dependent or threshold rule [62]. Future work could aim to elucidate whether pigeons employ such rules in determining predator location.

Given its ubiquity, collective detection via gaze following may be widespread across many species or contexts, providing a rapid, low-cost mechanism for sharing information and coordinating group responses. However, we suggest that this strategy is particularly beneficial in species lacking explicit alarm signals, and/or in situations where false alarms are especially costly; for example, when escape is energetically demanding due to large body size, or when unnecessary flight results in lost foraging opportunities in patchy or competitive environments. We acknowledge that there may be other explicit alarm signals that occur later in the detection timeline, such as wing sonation [63], which operate together with gaze following. Recent advances in markerless, computer vision–based posture tracking [64–66] now provide scalable, non-invasive tools for extending this research to wild populations, opening new possibilities for comparative studies under natural conditions.

This study integrated fine-scale tracking [45, 46], information transmission modeling [48–50], sensory ecology [35, 47], animal cognition [28, 37, 61], and sensory network analysis [20, 51, 52]. Our approach and findings reflect a growing trend across disciplines toward recognizing that cognitive and collective processes are mutually necessary to fully understand social phenomena [67, 68]. Our results demonstrate how individual cognition and group dynamics operate jointly to enhance the adaptive value of social living.

## Materials and Methods

### Ethics statement

All animal experiments were approved under license 35-9185.81/G-19/107 by the Regierungspräsidium Freiburg, the animal ethics authority of Baden-Württemberg, Germany. Pigeons were monitored daily and housed with perches and nesting structures. No pigeons were injured during the simulated predator events.

| Term | Definition |
| --- | --- |
| Foveation | Directing the fovea—associated with high-acuity vision—toward a visual target. |
| Target monitor | The monitor displaying a looming raptor shadow during the simulated predator attack. |
| Distractor monitor | An identical monitor that remains blank (does not display the predator stimulus) during the simulated predator attack. |
| Informed individual | An individual that has foveated on the target monitor (has “detected” the predator cue). |
| Uninformed individual | An individual that has not yet foveated on the target monitor |
| Cue | A behaviour displayed by an informed individual that may be used by an uninformed individual to detect the predator stimulus. |

Glossary of terms used in the study

### Subjects

Twenty pigeons (*Columba livia*) were tested in this study (13 females and 7 males), all juveniles (approximately 1 year old), originating from the same breeder. They were housed together in an outdoor aviary, provided with perches and fed grains once daily. On experimental days, they were fed only after the experiment was completed to ensure foraging motivation. Water and grit were provided *ad libitum*.

### Experimental design

The experiment took place in the SMART-BARN [45], a large-scale tracking facility (15 m length × 7 m width × 4 m height) hosted by the Max Planck Institute of Animal Behavior. In the centre of the experimental room, a 4.2 × 3.6 m area was covered in jute fabric and scattered with seeds, allowing a flock of 10 pigeons to forage collectively. In two opposite corners, a monitor (61.5 cm width × 37 cm height, WQHD [2560 × 1440], 144 Hz, G-MASTER GB2760QSU, Iiyama) was placed on a table to display a looming predator shadow stimulus, simulating an approaching predator. Hidden beneath the tables, a plastic model predator could be run across the room on a wire using a motorized pulley system (Wiral LITE kit, Wiral Technologies AS).

We used the motion-capture system of the SMART-BARN, equipped with 32 motion-capture cameras (12 Vero v2.2 and 20 Vantage 5, VICON), to track the pigeons’ fine-scale head and body movements. A Styrofoam plate equipped with four motion-capture markers (9 mm diameter, OptiTrack), arranged in a unique combination, was attached to each pigeon’s back to identify individuals. An additional four motion-capture markers (6.4 mm diameter) were attached to the head feathers to track head position and orientation.

Each individual participated in six trials, with each trial consisting of two simulated predator incursions (each pigeon hence experienced 12 predator presentations). For every trial, the group of 20 pigeons was divided into two pseudo-randomly generated flocks of 10, ensuring that none of the 12 flock compositions were repeated across trials. We arranged the composition so that each pigeon was grouped with every other pigeon at least once across the six trials. Each individual experienced a maximum of one test per day. This design resulted in a total of 24 simulated predator events.

At the start of each experimental session, the 10 pigeons were equipped with motion-capture markers, and four photographs of each individual’s head were taken simultaneously from different angles using four webcams, to later reconstruct the relative positions of the eyes and beak in relation to the markers. After calibrating the motion-capture system, the pigeons were released into the experimental room. Following a 2-minute acclimatization period, the experimenter scattered a mix of grains and grit in the foraging area and then hid behind a curtain. Once at least half of the pigeons had begun pecking, a free-feeding period of 2–5 minutes started, with the duration randomized to reduce predictability. At the end of the feeding period, the experimenter triggered the looming stimulus on one of the monitors (lasting approximately 10 seconds), immediately followed by the model predator crossing the room on the wire. This sequence constituted one predator presentation event. After at least half of the pigeons had resumed feeding and another free-feeding period had elapsed, a second predator event was presented on the opposite side of the room using the second monitor and model predator.

After the session, the motion-capture markers were removed from the pigeons’ heads and bodies, and the birds were returned to their aviary. The data analyzed here originates from the study described in Delacoux & Kano (2024) [47], which provides further details on the experimental setup, procedures, and data processing.

### Data processing

The motion capture data was exported from the motion capture software (Nexus version 2.14, VICON) as a CSV of 3D coordinates, then processed with custom MATLAB code (provided in [69]).

Using four images of the pigeons’ heads, a custom structure-from-motion pipeline reconstructed the positions of the eyes and beak relative to four motion-capture markers. By aligning these reconstructed head positions with the 3D coordinates from the motion-capture system, we defined each pigeon’s local head coordinate system—representing the location of objects and conspecifics from the bird’s perspective—for the entire trial.

The resulting data were then filtered using a custom pipeline (described in supporting information of [46] and [47]) to smooth trajectories and remove improbable movements. From the local head coordinate system, we reconstructed the foveal gaze vector based on known projection angles: 75° in azimuth and 0° in elevation. We did not consider binocular vision in this analysis, as previous research has shown that pigeons rely exclusively on their foveas when attending to a distant predator image [47].

Based on the motion capture 3D coordinates, we approximated objects of interest (monitors and other pigeons) as 3D volumes. The monitors were represented as a single sphere of 86.13 cm (monitor’s diagonal length = 71.77 cm + 20% margin). The pigeons were defined as 2 spheres representing the head (12 cm diameter) and the body (24 cm diameter). These objects were then reprojected onto the head local coordinate system of the pigeons. After considering occlusion from these objects and the pigeon’s blind field (40^*°*^ at the back of the head), we calculated the area an object covered in an individual’s visual field. Additionally, a pigeon was defined as “foveating” on an object if one of its gaze vectors intersected the object (within a 10^*°*^ error margin to account for eye movements and tracking error).

To characterize the detection of the looming predator stimulus, we refined the definition of foveation on the monitor to ensure that the first foveation was not a random gaze crossing [47]. Our thresholds are conservative and biologically meaningful. Specifically, we excluded foveations occurring earlier than 200 ms after the onset of the looming cue, based on pigeons’ typical reaction times [70], and also excluded foveations shorter than 300 ms based on typical duration of fixations in pigeons [46].

Based on the motion capture data, we extracted several features to build four different types of networks (S3 Table) and seven behavioural cues exhibited by the pigeons (S10 Table, S3 Fig).

### Social transmission analysis

We constructed Bayesian model of social transmissions to ask 1) whether social transmission was important for foveation on the location of the predator, and if so, what were the 2) information pathways and 3) relevant cues that were responsible for transmission. We used the network-based diffusion analysis (NBDA) framework [54, 71, 72], estimating a hazard rate for a target event as a non-homogeneous Poisson process that included a combination of a social transmission rate (*s*^*′*^) and an intrinsic (asocial) rate (*λ*_0_). The relative strength of social influence *(s)* was then calculated by dividing the posterior distributions where *s* = *s*^*′*^*/λ*_0_. We additionally calculated the percentage of events that occurred through social transmission (%ST, see S1 Text). Here, the event of interest was becoming informed about the location of the predator, defined as when a pigeon first foveated on the looming predator stimulus for longer than 300ms. Thus, *λ*_0_ is interpreted as the propensity to foveate on the predator independently of social information. *s*^*′*^ is interpreted as the influence that cues from informed social partners, defined by a network, has on a focal individual to then themselves foveate on the predator. *s* is thus the strength of this social influence per unit connection relative to the asocial rate. We used dynamic networks with dynamic transmission weights sampled at 1Hz, so the model accounted for the changing spatial configuration of pigeons and the variable emission of relevant cues within a trial. We accounted for effects of distance from the target monitor and time spent head-down while foraging as time-varying individual-level variables. Main text models are fitted with varying effects to account for inter-individual differences of *s*^*′*^ and *λ*_0_. To determine if repeated exposures to the predator were a confounding factor, we fit a model that included trial (per individual) as a statistical control (summarized in S11 Table). Trial was not estimated to significantly affect either social transmission or asocial detection (*β*_*trial*_ = 0.021[*−* 0.024, 0.064]), and we exclude it from the main-text models for simplicity.

Models were constructed and fitted with the STbayes R package, a Bayesian implementation of NBDA [48]. Parameter point estimates have been shown to be nearly identical between STbayes and the original NBDA package, although the STbayes package allows for improved estimation of uncertainty in terms of parameter values and model comparison metrics. See S1 Text for details of the models and S2 Text for details of priors. To test the results robustness to priors, we refit the best fitting model with priors that were overly skeptical of social transmission (*s ≈* 0.36) and found similar estimates (S12 Table). Priors for co-variates were weakly informative and skeptical of large effects. A power analysis demonstrated that our sample size (24 trials) was more than sufficient, as support for the correct model under even a small effect of social transmission was achieved with a design of 5 or more trials (see supplementary code).

Our analysis strategy was similar to previous multi-network NBDA analyses [73–75]. We first fit a set of constrained models that excluded the possibility of social transmission. We then fit unconstrained models using all combinations of networks and cues of interest. We fit multi-network models which allowed for *s*^*′*^ to be estimated for each network simultaneously, and compared these with single network models where *s*^*′*^ was constrained to 0 for all other networks. We performed model comparison using Pareto smoothed importance-sampling leave-one-out cross-validation (PSIS-LOO) to determine relative support for social transmission and relevant networks and cues (see S3 Text) [56]. We repeated this process for both the target monitor and distractor monitor.

We took several steps to prepare the data for the model. For each trial, we measured event times as the frame at which each bird first foveated at the looming predator stimulus, which we assume was the moment that they became informed of the predator’s location. For each frame, we measured the edge weights between each dyad of our 4 possible transmission networks. Visual field and foveation were necessarily directed networks, while voronoi and inverse distance were undirected networks. Visual field and inverse distance were continuous measures, and we binarized these to be comparable to voronoi and foveation networks, such that *a*_*ijt*_ *≥* median(*a*) was set to 1, otherwise 0. We did not use a maximum distance threshold, as conspecifics and stimuli were large enough to be visible from any position in the experimental room.

To assess sensitivity to our median threshold, we tested binarization at the 40th and 60th percentiles in addition to the median. The visual field network became degenerate (i.e. all zeroes) below the 34th percentile, so we did not test lower thresholds. Correlations between continuous and binary edges were similar across thresholds. Binary networks generally increased the distinction between inverse distance and visual field networks, although the median threshold did not maximize the distinction between the two networks (S4 Fig), and thus did not bias our results. We also confirmed that the 4 transmission networks were not overly correlated with one another (S5 Fig). Continuous distributions and thresholds are shown in S6 Fig. Main results were also robust to threshold choice (S7 Fig, S8 Fig, S9 Fig), and we present the 50th percentile in the main text.

For each frame, we also calculated the dynamic transmission weight for each cue, which was *w*_*jt*_ = 1 if the cue was being emitted by individual *j* at time *t*, and otherwise 0. This ensured the unit connection between individuals represented 1 frame of connection to an informed, cueing flock-mate.

We used the “high-resolution” feature of STbayes to fit the model. Rather than calculating a summary of *a*_*ijt*_ and *w*_*ijt*_ per inter-event interval, this value was calculated per frame. This accounted for cases when network connections and cue emission did not align, which would be possibly conflated. This process was done for each network *n* separately.

For the multi-network model, the hazard rate for bird *i* at frame *t* was defined as:

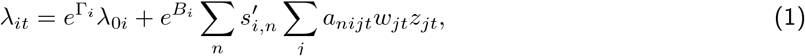

where the first term is the intrinsic rate, and the second term is the social rate. Γ contained co-variates affecting the intrinsic rate and *B*_*i*_ contained the co-variates affecting the social transmission rate. ILVs are exponentiated, such that a coefficient of 0 does not affect the overall rate of becoming informed. *λ*_0_ and 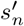 were fit with varying effects per individual. *z*_*jt*_ is a state variable that indicates whether *j* is already informed, such that only informed individuals influenced naive individuals. All models were fit with 5 chains with 2500 warmup iterations and 2500 sampling iterations. Models converged well, with chain mixing assessed visually from trace-plots and Rhat values for all parameters were *≤* 1.001. We verified that parameters were well identified for the best fitting main-text model by plotting the posterior versus prior distributions (S10 Fig). To verify the most-predictive model fit the data well, we additionally performed a posterior predictive check where diffusion curves for all trials were simulated using posterior distributions of parameters from the best-fitting single network model (S11 Fig).

## Acknowledgements

We thank all lab members, animal keepers, veterinarians, technicians, and administrative staff at the Centre for the Advanced Study of Collective Behaviour (CASCB) and Max-Planck Institute of Animal Behavior (MPI-AB). During the preparation of this work the author(s) used ChatGPT 5 in order to refine existing text for clarity and concision. After using this tool/service, the author(s) reviewed and edited the content as needed and take(s) full responsibility for the content of the published article.

## Authors contributions

Conceptualization: MD, MC, FK

Methodology: MD, MC, FK

Investigation: MD, MC, FK

Visualization: MD, MC

Funding Acquisition: MC, FK

Supervision: MC, FK

Writing - Original Draft: MD, MC, FK

Writing - Review & Editing: MD, MC, FK

## Data Availability

Code and data underlying statistical analyses and figures are available on Edmond via https://doi.org/10.17617/3.AWIUN8.

## Supporting information

### Text

- **S1 Text. Extended description of network-based diffusion analysis models**.
- **S2 Text. Description of choice of priors**.
- **S3 Text. Description of model comparison**.

## Figures

- **S1 Fig. Evolution of the number of pigeons looking at the target monitor at least one time over the course of the trials**. Trials are arranged in the order they occurred (x-axis). The number of individuals who detect the predator remained consistent over trials. The data underlying this Figure can be found in https://doi.org/10.17617/3.AWIUN8.
- **S2 Fig. Model comparison of “distractor” models**. We used LOO-PSIS to compare the models that predicted the distractor monitor detection transmission. Control analysis D tested if the pattern of foveations on the distractor monitor was better predicted by the informed flockmates foveating (“cueing”) on the distractor or the target monitor (dark blue). This analysis is conceptually similar to Control analysis C, where the “staring” behavior of informed flockmates would lead the uninformed individual to look at a monitor, though not the same one foveated by the informed individuals. In a supplementary control analysis, we tested a null model which did not include social information (detection only through individual learning; light green), as well as to the different cues of the flockmates (dark green). Similarly to the “target” models, the duration of foveation on the distractor monitor (main) model was best predicting the transmission. This suggests that pigeons followed the gaze of conspecifics towards the distractor monitor, despite the absence of displayed stimulus. The data underlying this Figure can be found in https://doi.org/10.17617/3.AWIUN8.
- **S3 Fig. Evolution of the different behaviors over the course of the trial**. The left graphs show the average number of individuals displaying a certain behavior at each given timepoint. The top graph represents the average number of individuals foveating on the target or the distractor, while the bottom graph shows the number of individuals in a head-up position, walking, running away from the target monitor or flying. The right graphs represent the average frequency of becoming head-up (top) and number of head-scan saccades (bottom). As these behaviors are point events rather than continuous behaviors, we represented the data as bar plots showing the number of occurrences within each 0.5-second interval of the trial. The data underlying this Figure can be found in https://doi.org/10.17617/3.AWIUN8.
- **S4 Fig. Correlations between binarized and continuous network edges of visual field and inverse distance networks at three thresholds**. For all thresholds, correlations between continuous and binary networks are stable across the a thresholds, suggesting that the median binarization did not introduce any artifacts. Furthermore, Binarization increased the distinction between avf and inverse distance networks, minimizing the information shared between them. The correlation between the two binary networks was lowest at the 40th percentile (0.27). The data underlying this Figure can be found in https://doi.org/10.17617/3.AWIUN8.
- **S5 Fig. Correlations of edges from all four networks**. Correlation plot for the binarized networks obtained from the target monitor dataset. Networks were not overly correlated. The data underlying this Figure can be found in https://doi.org/10.17617/3.AWIUN8.
- **S6 Fig. Distribution of continuous edge weights prior to binarization for visual field and inverse distance networks**. Continuous edge distributions shown for inverse distance (blue) and visual field (red) networks. Vertical lines indicate the three thresholds we tested for binarization (.4, .5, and .6). The data underlying this Figure can be found in https://doi.org/10.17617/3.AWIUN8.
- **S7 Fig. Estimated social transmission rates (***s*^*′*^**) from the best-fitting multi-network model (fov-mon) across binarization thresholds (40th, 50th, 60th percentile)**. Points show posterior medians; error bars show 95% credible intervals. Estimates were robust to the choice of threshold. The data underlying this Figure can be found in https://doi.org/10.17617/3.AWIUN8.
- **S8 Fig. Sensitivity of model comparison results to the 40th percentile binarization threshold for continuous network edges**. LOOIC scores shown for multi-network models with different transmission weight cues. The dashed line indicates the best model. The data underlying this Figure can be found in https://doi.org/10.17617/3.AWIUN8.
- **S9 Fig. Sensitivity of model comparison results to the 60th percentile binarization threshold for continuous network edges**. LOOIC scores shown for multi-network models with different transmission weight cues. The dashed line indicates the best model. The data underlying this Figure can be found in https://doi.org/10.17617/3.AWIUN8.
- **S10 Fig. Estimated posterior distributions versus prior distributions for the best-fitting model**. *log*(*s*^*′*^) indexes correspond to area visual field, distance, voronoi, and foveation networks. Parameters were well identified. The data underlying this Figure can be found in https://doi.org/10.17617/3.AWIUN8.
- **S11 Fig. Posterior predictive check**. We simulated diffusion curves for each trial from 1000 draws of the posterior (each thin line is 1 draw). Overall, the shape of diffusion curves predicted by the model match the actual data (thick lines). The data underlying this Figure can be found in https://doi.org/10.17617/3.AWIUN8.
- **S12 Fig. Pareto diagnostics for multi-network models (target monitor)**. Despite several observations receiving a score over 0.7, none received a score over 1.0. This indicated that PSIS-LOO evaluations were likely reliable. The data underlying this Figure can be found in https://doi.org/10.17617/3.AWIUN8.
- **S13 Fig. Pareto diagnostics for multi-network models (distractor monitor)** No observations received a score over 0.7. This indicated that PSIS-LOO evaluations were likely reliable. The data underlying this Figure can be found in https://doi.org/10.17617/3.AWIUN8.

## Tables

- **S1 Table. Model comparison for purely intrinsic models (asocial learning) for foveating on the target monitor**. We include each models LOOIC score, the standard error of the difference in LOOIC scores to the best fitting model, the difference in ELPD (expected log posterior density) to the best fitting model, and the standard error of that difference. Information related to the distance to the monitor, as well as the amount of time spent head down positively influenced predictive performance of models. The best LOOIC score was obtained when including the area of the visual field that the monitor took up, and the proportion of time spent head down as ILVi. The data underlying this Table can be found in https://doi.org/10.17617/3.AWIUN8.
- **S2 Table. Model comparison for purely intrinsic models (asocial learning) for foveating on the distractor monitor**. We include each models LOOIC score, the standard error of the difference in LOOIC scores to the best fitting model, the difference in ELPD (expected log posterior density) to the best fitting model, and the standard error of that difference. Similarly to the target monitor, information related to the distance to the monitor and the amount of time spent head down positively influenced predictive performance of models. The best LOOIC score was obtained when including the area of the visual field that the monitor took up, and the proportion of time spent head down as ILVi. The data underlying this Table can be found in https://doi.org/10.17617/3.AWIUN8.
- **S3 Table. Descriptions of networks used in the models**.
- **S4 Table. Posterior summary statistics for model parameters from best fitting multi-network model (“target” model)**. Columns include parameter name, median value, median absolute deviation (MAD), the lower and upper bounds of the 95% highest posterior density estimates, the number of effective samples and Rhat score. Parameters are fit on the log scale, and thus we included *log*(*λ*_0_) and *log*(*s*^*′*^), as well as their standard interpretations. *β* parameters are individual-level variables for the intrinsic rate (ILVi) and social rate (ILVs). *σ*_*ID*_ parameters are the scale parameter for varying effects. %*ST* gives the estimated percent of events that occurred through social transmission. The data underlying this Table can be found in https://doi.org/10.17617/3.AWIUN8.
- **S5 Table. Model comparison for multi-network and all single network models (“target” models)**. We include each models LOOIC score, the standard error of the difference in LOOIC scores to the best fitting model, the difference in ELPD (expected log posterior density) to the best fitting model, and the standard error of that difference. The data underlying this Table can be found in https://doi.org/10.17617/3.AWIUN8.
- **S6 Table. Posterior summary statistics for model parameters from best fitting single network (“target” model)**. Columns include parameter name, median value, median absolute deviation (MAD), the lower and upper bounds of the 95% highest posterior density estimates, the number of effective samples and Rhat score. Parameters are fit on the log scale, and thus we included *log*(*λ*_0_) and *log*(*s*^*′*^), as well as their standard interpretations. *β* parameters are individual-level variables for the intrinsic rate (ILVi) and social rate (ILVs). *σ*_*ID*_ parameters are the scale parameter for varying effects. The data underlying this Table can be found in https://doi.org/10.17617/3.AWIUN8.
- **S7 Table. Model comparison for multi-network and all single network models (“distractor” models)**. We include each models LOOIC score, the standard error of the difference in LOOIC scores to the best fitting model, the difference in ELPD (expected log posterior density) to the best fitting model, and the standard error of that difference. The data underlying this Table can be found in https://doi.org/10.17617/3.AWIUN8.
- **S8 Table. Posterior summary statistics for model parameters from best fitting multi-network model (“distractor” model)**. Columns include parameter name, median value, median absolute deviation (MAD), the lower and upper bounds of the 95% highest posterior density estimates, the number of effective samples and Rhat score. Parameters are fit on the log scale, and thus we included *log*(*λ*_0_) and *log*(*s*^*′*^), as well as their standard interpretations. *β* parameters are individual-level variables for the intrinsic rate (ILVi) and social rate (ILVs). *σ*_*ID*_ parameters are the scale parameter for varying effects. %*ST* gives the estimated percent of events that occurred through social transmission. The data underlying this Table can be found in https://doi.org/10.17617/3.AWIUN8.
- **S9 Table. Posterior summary statistics for model parameters from best fitting single network (“distractor” model)**. The data underlying this Table can be found in https://doi.org/10.17617/3.AWIUN8.
- **S10 Table. Description of the different behavioural cues used as alternatives for the foveation on the monitor**.
- **S11 Table. Posterior summary statistics for model parameters from multi-network model, refit with trial as a multiplicative effect**. Trial was not estimated to affect either asocial or social learning pathways. Columns include parameter name, median value, median absolute deviation (MAD), the lower and upper bounds of the 95% highest posterior density estimates, the number of effective samples and Rhat score. Parameters are fit on the log scale, and thus we included *log*(*λ*_0_) and *log*(*s*^*′*^), as well as their standard interpretations. *β* parameters are individual-level variables for the intrinsic rate (ILVi) and social rate (ILVs). *σ*_*ID*_ parameters are the scale parameter for varying effects. The data underlying this Table can be found in https://doi.org/10.17617/3.AWIUN8.
- **S12 Table. Posterior summary statistics for model parameters from best-fitting multi-network model, refit with a prior more skeptical of social transmission (***log*(*λ*_0_) *∼* ( *−*5, 2), *log*(*s*^*′*^) *∼* ( *−*5, 2), **thus** *s ≈* 0.36). Estimates are similar to priors that assume *s* = 1. Estimates are similar, but with additional shrinkage. Columns include parameter name, median value, median absolute deviation (MAD), the lower and upper bounds of the 95% highest posterior density estimates, the number of effective samples and Rhat score. Parameters are fit on the log scale, and thus we included *log*(*λ*_0_) and *log*(*s*^*′*^), as well as their standard interpretations. *β* parameters are individual-level variables for the intrinsic rate (ILVi) and social rate (ILVs). *σ*_*ID*_ parameters are the scale parameter for varying effects. The data underlying this Table can be found in https://doi.org/10.17617/3.AWIUN8.

### Video

- **S1 Video. Video abstract**.

