## Supporting text, figures and tables for "Pigeons rapidly transmit information about predator location through gaze-following"

### Supporting Information

#### S1 Extended description of network-based diffusion analysis models

We wished to understand 1) whether or not social transmission played a role in the detection of the looming predator stimulus, 2) whether visual observation of others was important for social transmission, and 3) which specific cues were responsible for social transmission of information about the looming predator stimulus. To address these three questions, we constructed a set of Bayesian models of social transmission that were similar in design to network-based diffusion analysis models [1–3]. NBDA models are widely used in the social learning and animal culture literature to determine whether social transmission of novel behaviors is occurring, how strong its effects are, and the exact pathways of information transfer [4].

Typically, model comparison can be used to compare between a) a purely asocial model that does not consider any socially observed information and b) a model that contains some information about the connections between individuals (e.g. association) and the knowledge state of individuals (naive or informed). The asocial model estimates an intrinsic rate ($\lambda_0$), as well as the effect of any individual-level co-variates (ILVs) on the intrinsic rate. In our case, we believed apriori that proximity to the monitor where the stimulus would be displayed and whether or not birds were head-down (i.e. foraging) would affect the intrinsic rate, and thus included them as ILVs.

The social model estimates an intrinsic rate and a social transmission rate ($s'$) that affects the overall rates of an event occurring per unit connection. The relationship between $s'$ and $\lambda_0$, $s = s'/\lambda_0$, defines the strength of social transmission. If $s'$ is relatively small compared to $\lambda_0$, social transmission is likely not occurring, and vice versa. If $s' = \lambda_0$, this indicates that both asocial and social learning are important for the target event. Similarly, ILVs can be applied to $s'$. We apriori believed that the time spent in a head-down position would also negatively impact $s'$, and thus this was included as an ILV. Coefficients for the time head-down on intrinsic and social rates were estimated independently.

The standard NBDA model cannot distinguish which networks are most important for information transfer, but it can be extended to estimate different $s$ parameters for each network [5–7]. Within a multi-network model, if $s$ for a particular network is found to be near zero, this suggests that it may not be an important pathway of social transmission. Model comparison between the full multi-network model and models where $s'$ is constrained to be 0 then can be used to establish whether or not the inclusion of these networks improves predictive fit [3].

We additionally calculated the percentage of events that occurred through social transmission (%ST) on each network in our multi-network models. The probability that a single event occurs through social transmission on a given network is

$$P(\text{e occurred through ST on network n at time t}) = \frac{e^{\Gamma_i} s' \sum_j a_{ijt} z_{jt} w_{jt}}{\lambda_{t_e}}, \quad (1)$$

The overall percentage is then the average of these probabilities per network [3].

#### S2 Description of choice of priors

We mostly followed the default priors of the STbayes package, which are justified using prior predictive checks in Chimento and Hoppitt [8]. These priors allow for a wide range of plausible values, and assume apriori that rate of social transmission is no stronger than the intrinsic rate (i.e. $s = 1$). The priors are as follows:

$$\log(\lambda_0) \sim N(-4,2), \quad (2)$$
$$\log(s') \sim N(-4,2), \quad (3)$$
$$B_M \sim N(0,1), \quad (4)$$
$$B_{HD1} \sim N(0,1), \quad (5)$$
$$B_{HD2} \sim N(0,1), \quad (6)$$
$$z_{ID} \sim N(0,1), \quad (7)$$
$$\sigma_{ID} \sim \text{half}N(0,.5), \quad (8)$$
$$\rho_{ID} \sim \text{lkj corr cholesky}(4) \quad (9)$$
$$(10)$$

We applied more conservative prior of $N(0,.5)$ to the standard deviation of varying effects to prevent divergent transitions due to parameter identifiability. The prior of $\rho_{ID} \sim$ lkj corr cholesky$(4)$ indicates a prior belief that the covariance structure of varying-effects parameters are more likely to be independent.

### S3 Description of model comparison

We used the loo package to compute Pareto smoothed importance-sampling leave-one-out cross-validation (PSIS-LOO) for model comparison and report LOOIC [9]. LOOIC measures out-of-sample predictive performance—lower scores are better. We computed pairwise differences in LOOIC between the best fitting model and all other models in the set. If the standard error of the pairwise difference did not cross zero, we considered those LOOIC scores to be significantly different. Pareto diagnostics (visualized in figures S12, S13) indicated that some models contained 1 observation with $k > .7$, although none of these exceeded $k > 1$. This was not common enough for LOO estimates to be unreliable. "Target" models and "distractor" models were compared separately as they were fit to different data sets. We emphasize their LOOIC scores are not comparable.

#### S4 Supporting Figures

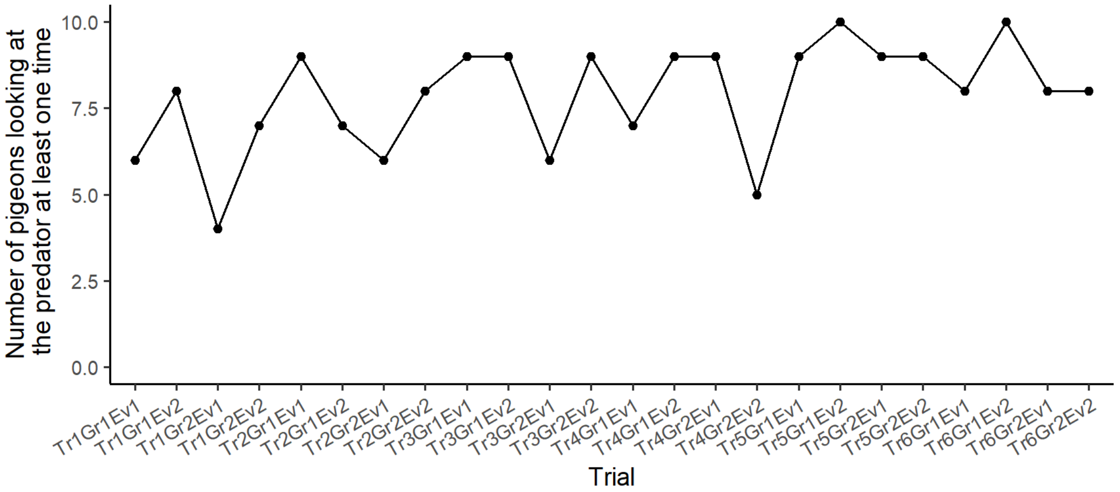


Figure S1: Evolution of the number of pigeons looking at the target monitor at least one time over the course of the trials. Trials are arranged in the order they occurred (x-axis). The number of individuals who detect the predator remained consistent over trials.

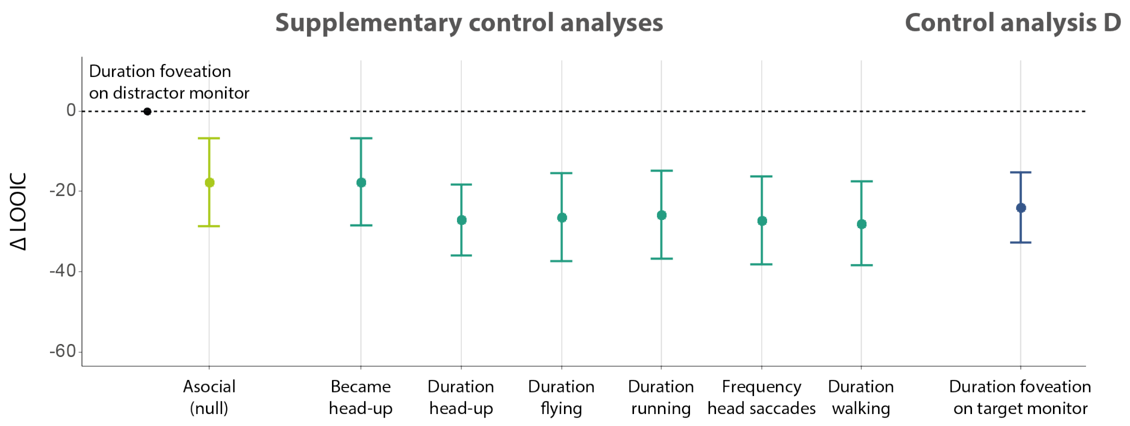


Figure S2: Model comparison of "distractor" models. We used LOO-PSIS to compare the models that predicted the distractor monitor detection transmission. Control analysis D tested if the pattern of foveations on the distractor monitor was better predicted by the informed flockmates foveating ("cueing") on the distractor or the target monitor (dark blue). This analysis is conceptually similar to Control analysis C, where the "staring" behavior of informed flockmates would lead the uninformed individual to look at a monitor, though not the same one foveated by the informed individuals. In a supplementary control analysis, we tested a null model which did not include social information (detection only through individual learning; light green), as well as to the different cues of the flockmates (dark green). Similarly to the "target" models, the duration of foveation on the distractor monitor (main) model was best predicting the transmission. This suggests that pigeons followed the gaze of conspecifics towards the distractor monitor, despite the absence of displayed stimulus.

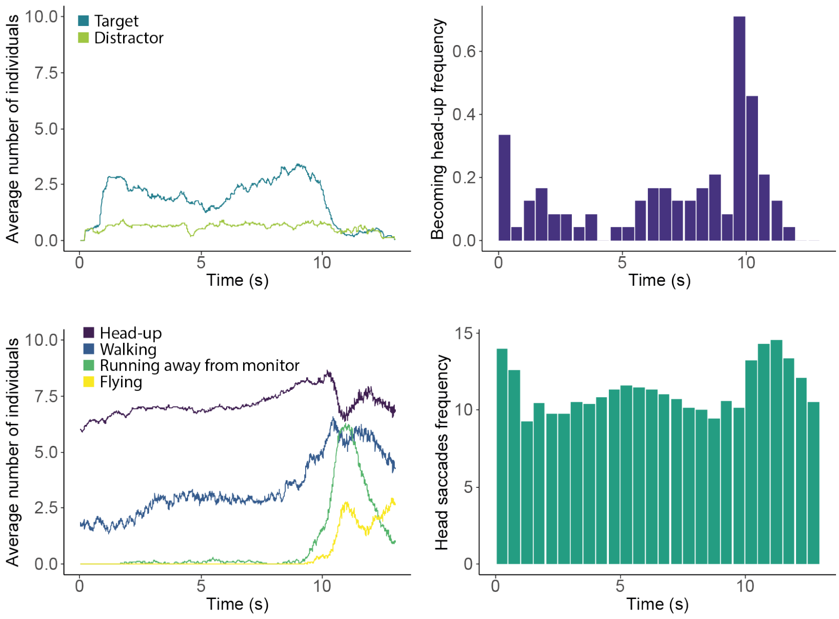


Figure S3: Evolution of the different behaviors over the course of the trial. The left graphs show the average number of individuals displaying a certain behavior at each given timepoint. The top graph represents the average number of individuals foveating on the target or the distractor, while the bottom graph shows the number of individuals in a head-up position, walking, running away from the target monitor or flying.The right graphs represent the average frequency of becoming head-up (top) and number of head-scan saccades (bottom). As these behaviors are point events rather than continuous behaviors, we represented the data as bar plots showing the number of occurrences within each 0.5-second interval of the trial.

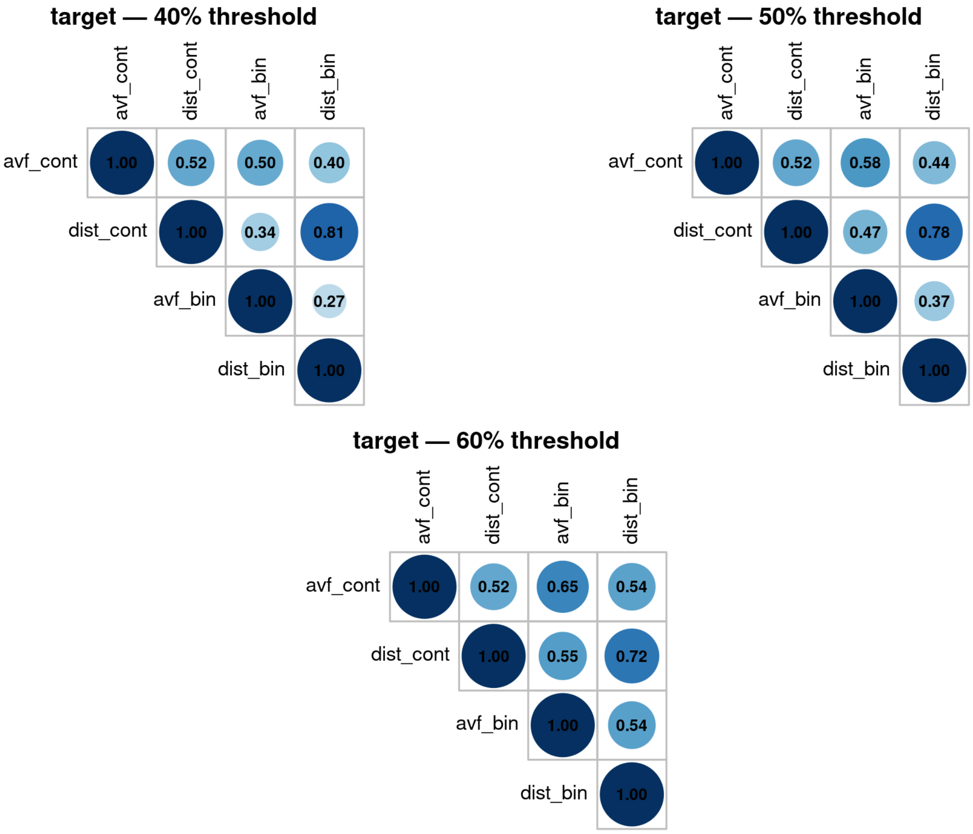


Figure S4: Correlations between binarized and continuous network edges of visual field and inverse distance networks at three thresholds. For all thresholds, correlations between continuous and binary networks are stable across the a thresholds, suggesting that the median binarization did not introduce any artifacts. Furthermore, Binarization increased the distinction between avf and inverse distance networks, minimizing the information shared between them. The correlation between the two binary networks was lowest at the 40th percentile (0.27).

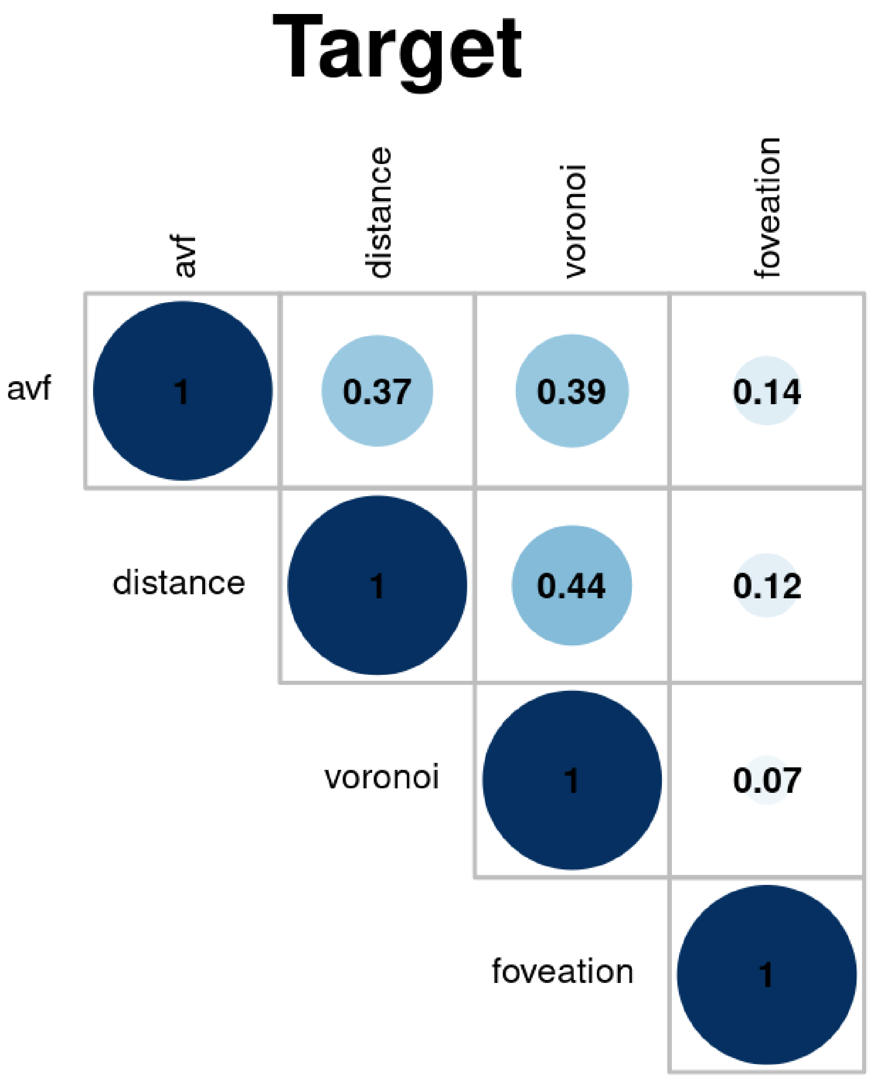


Figure S5: Correlations of edges from all four networks. Correlation plot for the binarized networks obtained from the target monitor dataset. Networks were not overly correlated.

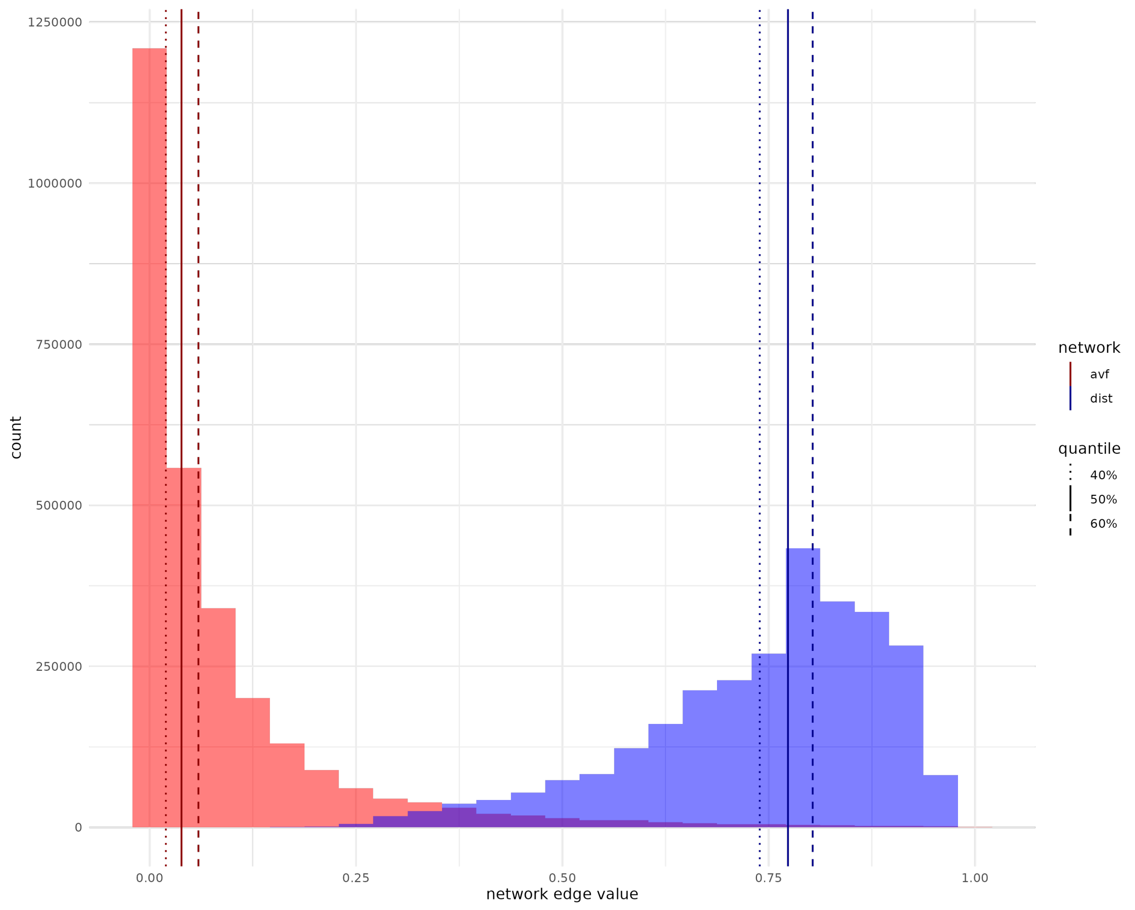


Figure S6: Distribution of continuous edge weights prior to binarization for visual field and inverse distance networks. Continuous edge distributions shown for inverse distance (blue) and visual field (red) networks. Vertical lines indicate the three thresholds we tested for binarization (.4, .5, and .6).

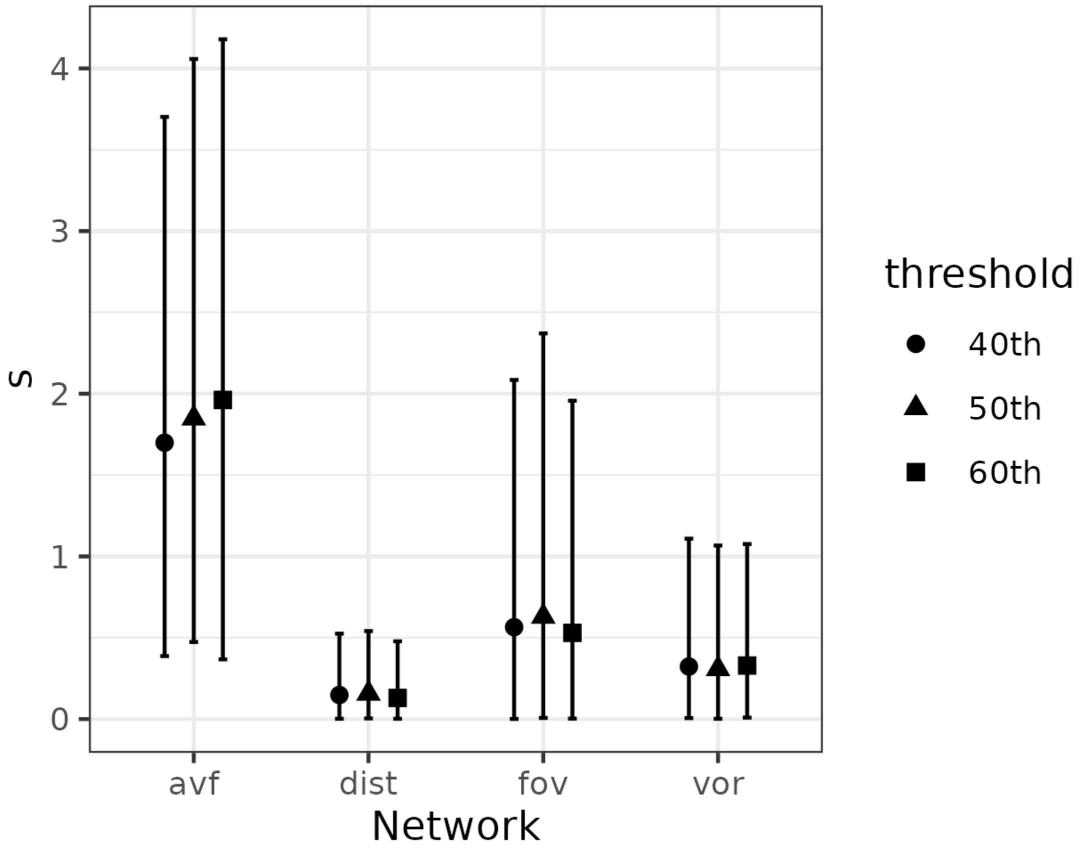


Figure S7: Estimated social transmission rates ($s'$) from the best-fitting multi-network model (fovmon) across binarization thresholds (40th, 50th, 60th percentile). Points show posterior medians; error bars show 95% credible intervals. Estimates were robust to the choice of threshold.

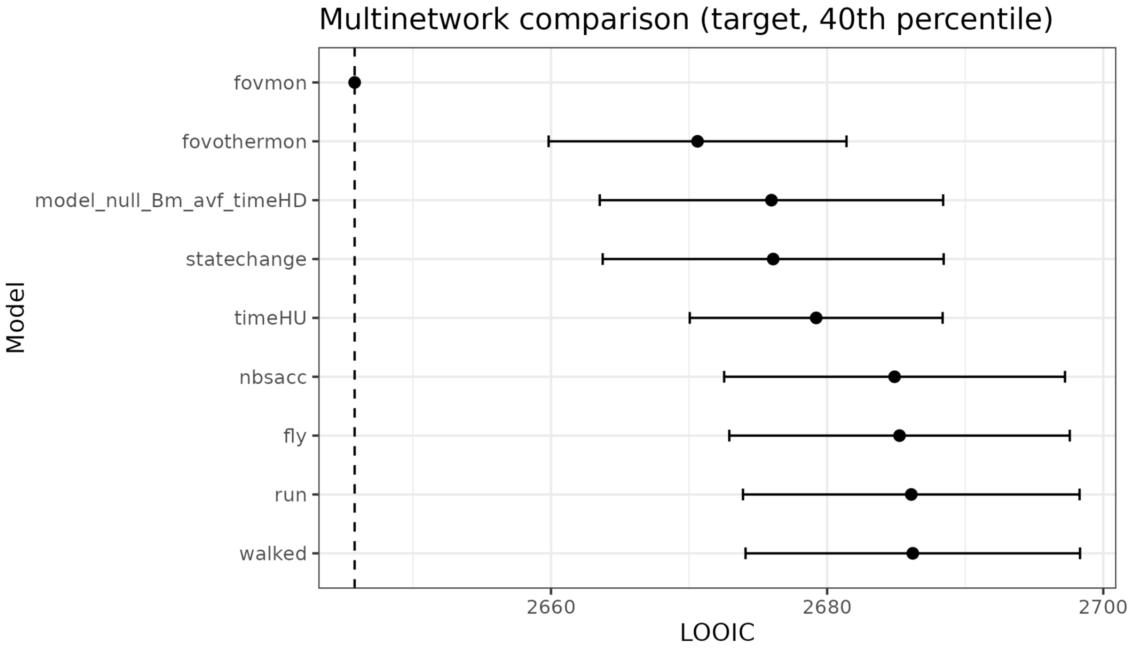


Figure S8: Sensitivity of model comparison results to the 40th percentile binarization threshold for continuous network edges. LOOIC scores shown for multi-network models with different transmission weight cues. The dashed line indicates the best model.

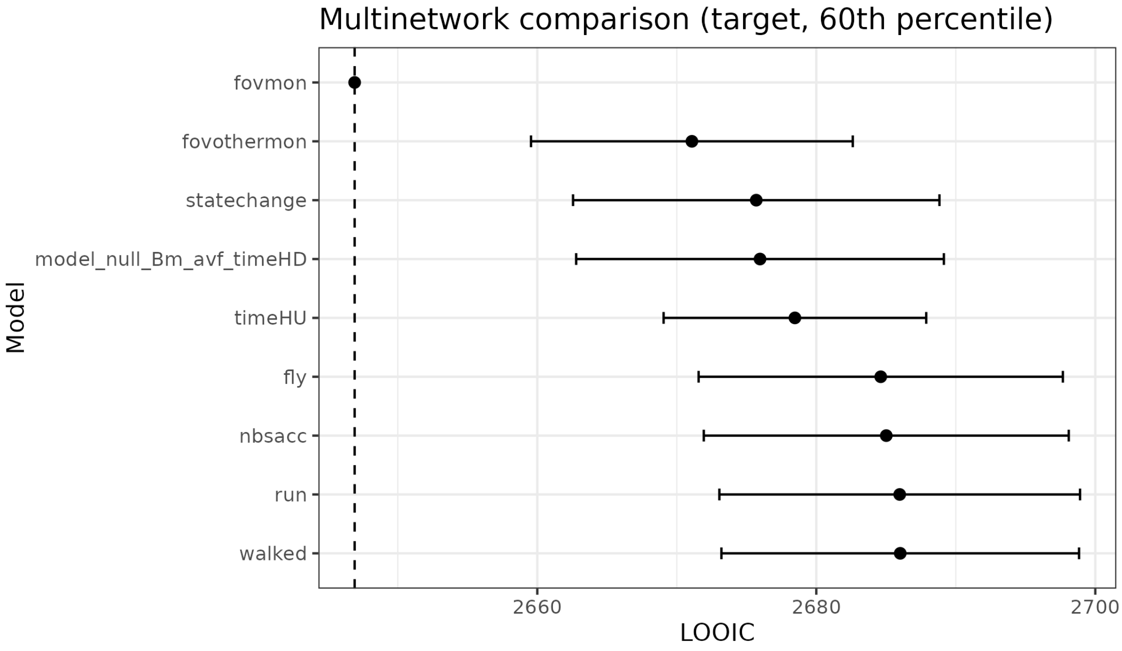


Figure S9: Sensitivity of model comparison results to the 60th percentile binarization threshold for continuous network edges. LOOIC scores shown for multi-network models with different transmission weight cues. The dashed line indicates the best model.

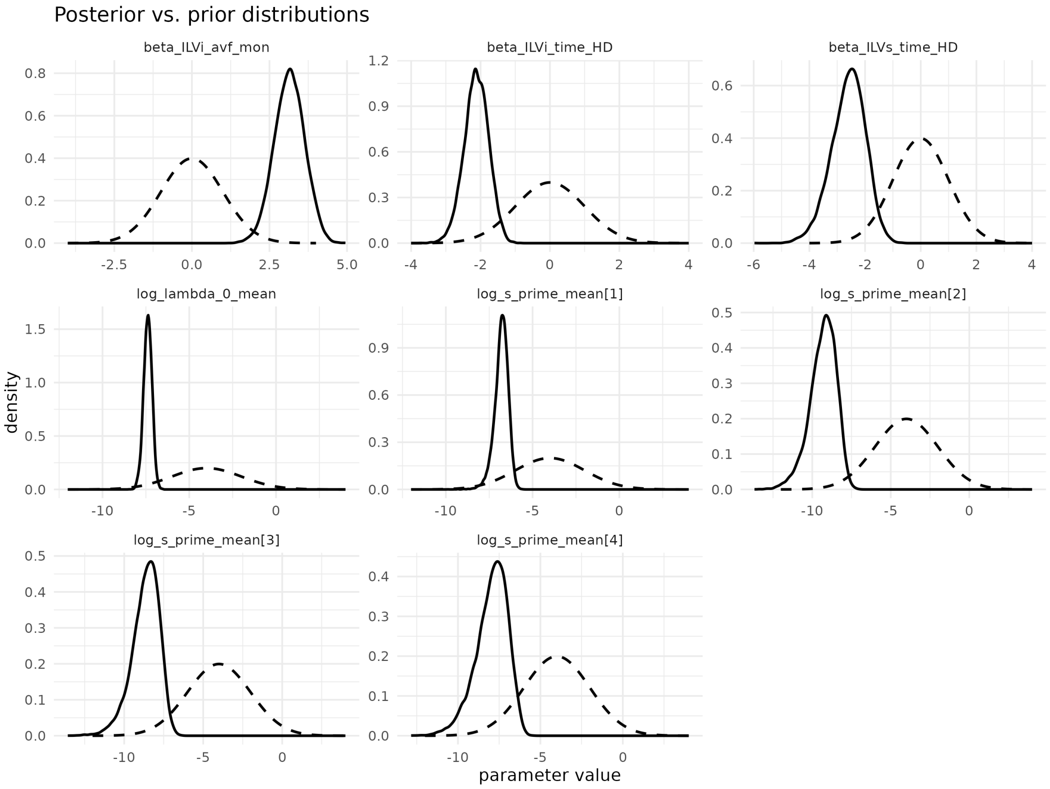


Figure S10: Estimated posterior distributions versus prior distributions for the best-fitting model. $log(s')$ indexes correspond to area visual field, distance, voronoi, and foveation networks. Parameters were well identified.

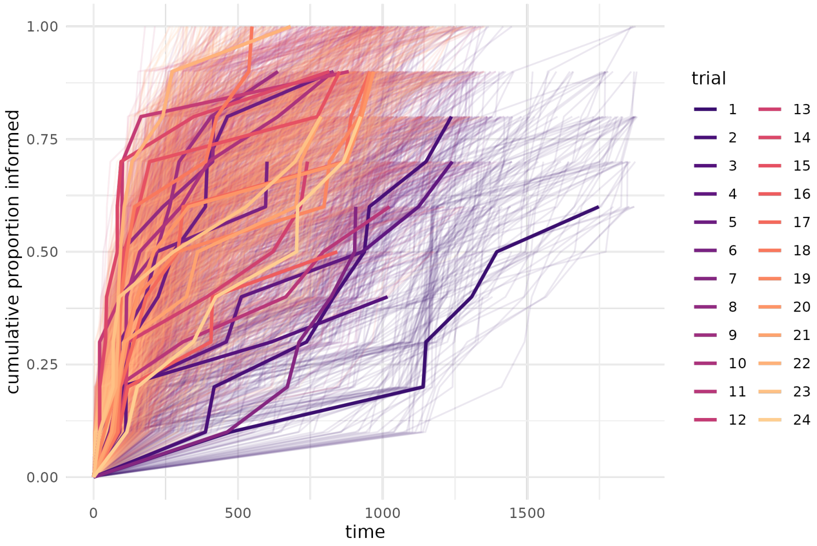


Figure S11: Posterior predictive check. We simulated diffusion curves for each trial from 1000 draws of the posterior (each thin line is 1 draw). Overall, the shape of diffusion curves predicted by the model match the actual data (thick lines).

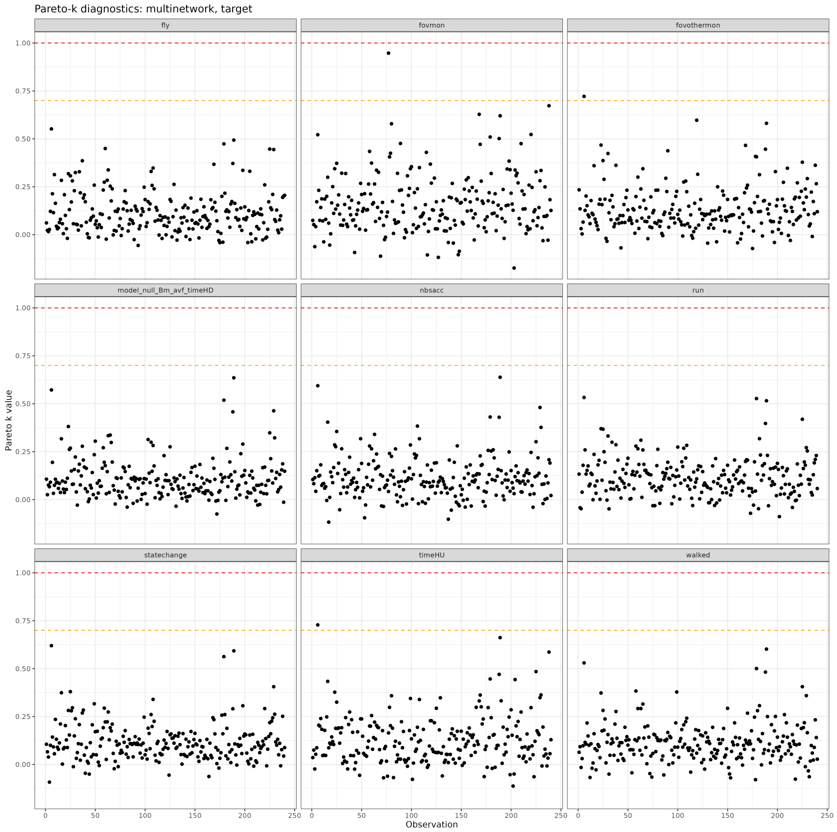


Figure S12: Pareto diagnostics for multi-network models (target monitor). Despite several observations receiving a score over 0.7, none received a score over 1.0. This indicated that PSIS-LOO evaluations were likely reliable.

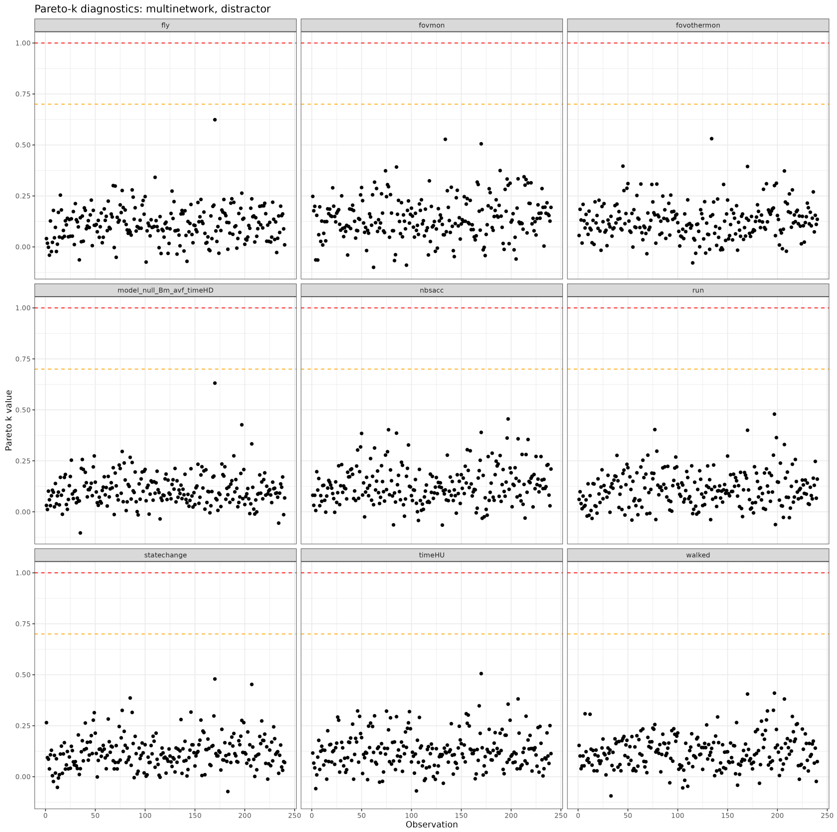


Figure S13: Pareto diagnostics for multi-network models (distractor monitor). No observations received a score over 0.7. This indicated that PSIS-LOO evaluations were likely reliable.

### S5 Supporting Tables

| Model_name | looic | se_looic_diff | elpd_diff | se_diff |
|---|---|---|---|---|
| ILVi: mon. avf, time HD | 2675.966 | 0.000 | 0.000 | 0.000 |
| ILVi: mon. dist, time HD | 2738.654 | 12.680 | -31.344 | 6.340 |
| ILVi: time HD | 2739.542 | 14.186 | -31.788 | 7.093 |
| ILVi: mon. avf | 2811.667 | 22.425 | -67.850 | 11.212 |
| ILVi: mon. dist | 2873.187 | 25.062 | -98.610 | 12.531 |
| No ILVi | 2878.878 | 26.105 | -101.456 | 13.052 |

Table S1: Model comparison for purely intrinsic models (asocial learning) for foveating on the target monitor. We include each models LOOIC score, the standard error of the difference in LOOIC scores to the best fitting model, the difference in ELPD (expected log posterior density) to the best fitting model, and the standard error of that difference. Information related to the distance to the monitor, as well as the amount of time spent head down positively influenced predictive performance of models. The best LOOIC score was obtained when including the area of the visual field that the monitor took up, and the proportion of time spent head down as ILVi.

| Model_name | looic | se_looic_diff | elpd_diff | se_diff |
|---|---|---|---|---|
| ILVi: mon. avf, time HD | 2216.466 | 0.000 | 0.000 | 0.000 |
| ILVi: time HD | 2231.068 | 6.949 | -7.301 | 3.474 |
| ILVi: mon. dist, time HD | 2232.539 | 6.537 | -8.037 | 3.269 |
| ILVi: mon. avf | 2277.808 | 14.673 | -30.671 | 7.336 |
| No ILVi | 2291.339 | 15.930 | -37.436 | 7.965 |
| ILVi: mon. dist | 2292.857 | 15.922 | -38.196 | 7.961 |

Table S2: Model comparison for purely intrinsic models (asocial learning) for foveating on the distractor monitor. We include each models LOOIC score, the standard error of the difference in LOOIC scores to the best fitting model, the difference in ELPD (expected log posterior density) to the best fitting model, and the standard error of that difference. Similarly to the target monitor, information related to the distance to the monitor and the amount of time spent head down positively influenced predictive performance of models. The best LOOIC score was obtained when including the area of the visual field that the monitor took up, and the proportion of time spent head down as ILVi.

| Network | Type | Description |
|---|---|---|
| Distance | Undirected | The metric distance between individuals |
| Voronoi | Undirected | Two individuals are connected if they are neighbors in the Voronoi tessellation, determined via Delaunay triangulation |
| Foveation | Directed | An individual is connected to another if it is directly foveating on that individual |
| Visual field | Directed | The extent to which individuals are visible in each other's visual fields (as a measure of how "visible" they are to each other) |

Table S3: Descriptions of networks used in models.

| | Parameter | Median | MAD | HPDI Lower | HPDI Upper | ESS bulk | ESS tail | Rhat |
|---|---|---|---|---|---|---|---|---|
| 1 | $\log\lambda_0$ | -7.384 | 0.242 | -7.863 | -6.907 | 14010.610 | 9582.883 | 1.000 |
| 6 | $\log s'_{avf}$ | -6.767 | 0.362 | -7.578 | -6.049 | 7754.070 | 7039.172 | 1.001 |
| 7 | $\log s'_{dist}$ | -9.235 | 0.824 | -11.057 | -7.805 | 14401.232 | 7870.655 | 1.000 |
| 8 | $\log s'_{vor}$ | -8.553 | 0.834 | -10.374 | -7.036 | 10241.418 | 8633.154 | 1.000 |
| 9 | $\log s'_{fov}$ | -7.836 | 0.926 | -9.885 | -6.219 | 15554.599 | 8516.789 | 1.001 |
| 5 | $\lambda_0$ | 0.001 | 0.000 | 0.000 | 0.001 | 14010.452 | 9582.883 | 1.000 |
| 10 | $s_{avf}$ | 1.849 | 0.870 | 0.474 | 4.058 | 8024.604 | 7532.824 | 1.001 |
| 11 | $s_{dist}$ | 0.157 | 0.129 | 0.004 | 0.541 | 14079.356 | 7893.282 | 1.000 |
| 12 | $s_{vor}$ | 0.308 | 0.252 | 0.002 | 1.067 | 11028.606 | 9145.821 | 1.000 |
| 13 | $s_{fov}$ | 0.630 | 0.566 | 0.007 | 2.371 | 15364.637 | 8421.211 | 1.000 |
| 2 | $\beta_{\text{ILViavfmon}}$ | 3.149 | 0.490 | 2.218 | 4.130 | 18732.688 | 9192.680 | 1.000 |
| 3 | $\beta_{\text{ILVitimeHD}}$ | -2.117 | 0.348 | -2.840 | -1.453 | 18203.679 | 10145.076 | 1.000 |
| 4 | $\beta_{\text{ILVstimeHD}}$ | -2.562 | 0.605 | -3.894 | -1.489 | 16699.764 | 9754.134 | 1.001 |
| 14 | $\sigma_{ID,log(s'_{avf})}$ | 0.749 | 0.296 | 0.028 | 1.250 | 4554.725 | 4480.587 | 1.001 |
| 15 | $\sigma_{ID,log(s'_{dist})}$ | 0.329 | 0.292 | 0.000 | 0.963 | 12332.621 | 7822.913 | 1.001 |
| 16 | $\sigma_{ID,log(s'_{vor})}$ | 0.375 | 0.318 | 0.000 | 1.008 | 9561.083 | 7009.976 | 1.000 |
| 17 | $\sigma_{ID,log(s'_{fov})}$ | 0.324 | 0.281 | 0.000 | 0.921 | 12397.066 | 7074.496 | 1.000 |
| 18 | $\sigma_{ID,log(\lambda_0)}$ | 0.376 | 0.174 | 0.000 | 0.680 | 4501.412 | 4884.218 | 1.000 |
| 19 | $\%ST$ avf | 0.281 | 0.059 | 0.155 | 0.395 | 6004.932 | 6413.691 | 1.001 |
| 20 | $\%ST$ dist | 0.026 | 0.019 | 0.001 | 0.075 | 13485.465 | 8567.331 | 1.000 |
| 21 | $\%ST$ vor | 0.046 | 0.034 | 0.000 | 0.128 | 8878.545 | 8017.852 | 1.001 |
| 22 | $\%ST$ fov | 0.018 | 0.015 | 0.000 | 0.057 | 15259.127 | 8448.886 | 1.000 |

Table S4: Posterior summary statistics for model parameters from best fitting multi-network model ("target" model). Columns include parameter name, median value, median absolute deviation (MAD), the lower and upper bounds of the 95% highest posterior density estimates, the number of effective samples and Rhat score. Parameters are fit on the log scale, and thus we included $log(\lambda_0)$ and $log(s')$, as well as their standard interpretations. $\beta$ parameters are individual-level variables for the intrinsic rate (ILVi) and social rate (ILVs). $\sigma_{ID}$ parameters are the scale parameter for varying effects. $\%ST$ gives the estimated percent of events that occurred through social transmission.

| Model name | looic | se looic diff | elpd diff | se diff |
|---|---|---|---|---|
| avf fovmon | 2640.632 | 0.000 | 0.000 | 0.000 |
| multinetwork fovmon | 2644.794 | 4.039 | -2.081 | 2.019 |
| vor fovmon | 2659.378 | 10.000 | -9.373 | 5.000 |
| dist fovmon | 2668.304 | 10.115 | -13.836 | 5.057 |
| fov fovmon | 2670.310 | 10.831 | -14.839 | 5.415 |
| avf fovothermon | 2671.011 | 11.075 | -15.189 | 5.538 |
| vor fovothermon | 2672.250 | 11.850 | -15.809 | 5.925 |
| multinetwork fovothermon | 2673.341 | 11.345 | -16.354 | 5.673 |
| dist fovothermon | 2673.655 | 11.737 | -16.511 | 5.869 |
| fov fovothermon | 2675.732 | 12.200 | -17.550 | 6.100 |
| avf statechange | 2675.837 | 12.478 | -17.603 | 6.239 |
| intrinsic only (null) | 2675.966 | 12.517 | -17.667 | 6.258 |
| multinetwork statechange | 2676.060 | 12.492 | -17.714 | 6.246 |
| vor statechange | 2676.214 | 12.491 | -17.791 | 6.245 |
| dist statechange | 2676.231 | 12.505 | -17.799 | 6.252 |
| fov statechange | 2676.393 | 12.478 | -17.880 | 6.239 |
| avf timeHU | 2677.123 | 11.143 | -18.245 | 5.572 |
| fov fly | 2677.158 | 12.481 | -18.263 | 6.240 |
| fov run | 2677.774 | 12.440 | -18.571 | 6.220 |
| fov timeHU | 2677.791 | 11.912 | -18.579 | 5.956 |
| fov nbsacc | 2678.161 | 12.477 | -18.764 | 6.239 |
| fov walked | 2678.314 | 12.460 | -18.841 | 6.230 |
| vor timeHU | 2678.315 | 12.030 | -18.841 | 6.015 |
| dist timeHU | 2678.418 | 12.059 | -18.893 | 6.029 |
| avf fly | 2678.613 | 12.466 | -18.991 | 6.233 |
| vor run | 2678.689 | 12.464 | -19.028 | 6.232 |
| dist run | 2678.726 | 12.438 | -19.047 | 6.219 |
| vor fly | 2678.739 | 12.503 | -19.053 | 6.251 |
| dist fly | 2678.813 | 12.470 | -19.090 | 6.235 |
| dist nbsacc | 2678.842 | 12.453 | -19.105 | 6.227 |
| vor walked | 2678.908 | 12.349 | -19.138 | 6.174 |
| avf nbsacc | 2678.929 | 12.425 | -19.148 | 6.213 |
| avf run | 2678.999 | 12.419 | -19.184 | 6.210 |
| avf walked | 2679.090 | 12.294 | -19.229 | 6.147 |
| dist walked | 2679.118 | 12.380 | -19.243 | 6.190 |
| vor nbsacc | 2679.242 | 12.497 | -19.305 | 6.249 |
| multinetwork timeHU | 2683.142 | 10.499 | -21.255 | 5.250 |
| multinetwork fly | 2685.023 | 12.542 | -22.196 | 6.271 |
| multinetwork run | 2685.774 | 12.440 | -22.571 | 6.220 |
| multinetwork nbsacc | 2686.445 | 12.463 | -22.907 | 6.232 |
| multinetwork walked | 2686.542 | 12.400 | -22.955 | 6.200 |

Table S5: Model comparison for multi-network and all single network models ("target" models). We include each models LOOIC score, the standard error of the difference in LOOIC scores to the best fitting model, the difference in ELPD (expected log posterior density) to the best fitting model, and the standard error of that difference.

| | Parameter | Median | MAD | HPDI Lower | HPDI Upper | ESS bulk | ESS tail | Rhat |
|---|---|---|---|---|---|---|---|---|
| 1 | $\log\lambda_0$ | -7.264 | 0.228 | -7.721 | -6.824 | 7782.296 | 8755.749 | 1.001 |
| 2 | $\log s'_{avf}$ | -6.497 | 0.284 | -7.099 | -5.943 | 6723.491 | 8582.565 | 1.000 |
| 8 | $\lambda_0$ | 0.001 | 0.000 | 0.000 | 0.001 | 7782.266 | 8755.749 | 1.000 |
| 7 | $s_{avf}$ | 2.151 | 0.865 | 0.803 | 4.285 | 6689.463 | 8049.177 | 1.000 |
| 3 | $\beta_{\text{ILViavfmon}}$ | 2.987 | 0.473 | 2.052 | 3.937 | 9780.141 | 9135.286 | 1.000 |
| 4 | $\beta_{\text{ILVitimeHD}}$ | -2.231 | 0.357 | -2.958 | -1.558 | 10901.501 | 9200.694 | 1.000 |
| 5 | $\beta_{\text{ILVstimeHD}}$ | -2.366 | 0.583 | -3.619 | -1.295 | 10263.573 | 8048.618 | 1.000 |
| 9 | $\sigma_{ID,log(s'_{avf})}$ | 0.694 | 0.244 | 0.152 | 1.186 | 3897.941 | 2600.305 | 1.000 |
| 10 | $\sigma_{ID,log(\lambda_0)}$ | 0.349 | 0.172 | 0.001 | 0.646 | 3733.928 | 3531.183 | 1.000 |
| 11 | $\%ST$ avf | 0.358 | 0.046 | 0.264 | 0.448 | 7445.816 | 8240.460 | 1.000 |

Table S6: Posterior summary statistics for model parameters from best fitting single network ("target" model). Columns include parameter name, median value, median absolute deviation (MAD), the lower and upper bounds of the 95% highest posterior density estimates, the number of effective samples and Rhat score. Parameters are fit on the log scale, and thus we included $log(\lambda_0)$ and $log(s')$, as well as their standard interpretations. $\beta$ parameters are individual-level variables for the intrinsic rate (ILVi) and social rate (ILVs). $\sigma_{ID}$ parameters are the scale parameter for varying effects.

| Model name | looic | se looic diff | elpd diff | se diff |
|---|---|---|---|---|
| multinetwork fovmon | 2198.716 | 0.000 | 0.000 | 0.000 |
| avf fovmon | 2199.976 | 5.359 | -0.630 | 2.680 |
| vor fovmon | 2202.916 | 4.937 | -2.100 | 2.468 |
| dist fovmon | 2203.621 | 5.200 | -2.452 | 2.600 |
| fov fovmon | 2206.480 | 8.184 | -3.882 | 4.092 |
| fov statechange | 2216.236 | 10.913 | -8.760 | 5.457 |
| vor statechange | 2216.270 | 10.911 | -8.777 | 5.455 |
| multinetwork statechange | 2216.325 | 10.917 | -8.804 | 5.459 |
| dist statechange | 2216.411 | 10.916 | -8.847 | 5.458 |
| avf statechange | 2216.457 | 10.902 | -8.870 | 5.451 |
| intrinsic only (null) | 2216.466 | 10.967 | -8.875 | 5.483 |
| fov timeHU | 2217.150 | 10.058 | -9.217 | 5.029 |
| fov fly | 2217.915 | 10.912 | -9.599 | 5.456 |
| fov fovothermon | 2218.058 | 10.223 | -9.671 | 5.111 |
| fov run | 2218.095 | 10.859 | -9.689 | 5.429 |
| avf fovothermon | 2218.110 | 9.855 | -9.697 | 4.927 |
| fov nbsacc | 2218.205 | 10.919 | -9.744 | 5.460 |
| dist fovothermon | 2218.479 | 9.759 | -9.882 | 4.880 |
| vor fovothermon | 2218.502 | 10.055 | -9.893 | 5.027 |
| fov walked | 2218.567 | 10.810 | -9.925 | 5.405 |
| avf run | 2218.835 | 10.856 | -10.059 | 5.428 |
| dist run | 2218.901 | 10.879 | -10.092 | 5.439 |
| vor run | 2218.963 | 10.864 | -10.123 | 5.432 |
| avf fly | 2218.972 | 10.869 | -10.128 | 5.435 |
| avf timeHU | 2219.016 | 10.134 | -10.150 | 5.067 |
| dist fly | 2219.046 | 10.879 | -10.165 | 5.439 |
| vor fly | 2219.113 | 10.868 | -10.198 | 5.434 |
| avf nbsacc | 2219.383 | 10.849 | -10.333 | 5.425 |
| dist walked | 2219.406 | 10.693 | -10.345 | 5.347 |
| vor timeHU | 2219.419 | 10.294 | -10.351 | 5.147 |
| vor nbsacc | 2219.427 | 10.858 | -10.355 | 5.429 |
| dist nbsacc | 2219.429 | 10.841 | -10.356 | 5.421 |
| dist timeHU | 2219.530 | 10.210 | -10.407 | 5.105 |
| avf walked | 2219.608 | 10.688 | -10.446 | 5.344 |
| vor walked | 2219.655 | 10.708 | -10.470 | 5.354 |
| multinetwork fovothermon | 2222.769 | 8.692 | -12.026 | 4.346 |
| multinetwork run | 2224.460 | 10.975 | -12.872 | 5.488 |
| multinetwork fly | 2225.138 | 10.917 | -13.211 | 5.458 |
| multinetwork timeHU | 2225.813 | 8.849 | -13.548 | 4.425 |
| multinetwork nbsacc | 2225.918 | 10.925 | -13.601 | 5.463 |
| multinetwork walked | 2226.689 | 10.457 | -13.986 | 5.229 |

Table S7: Model comparison for multi-network and all single network models ("distractor" models). We include each models LOOIC score, the standard error of the difference in LOOIC scores to the best fitting model, the difference in ELPD (expected log posterior density) to the best fitting model, and the standard error of that difference.

| | Parameter | Median | MAD | HPDI Lower | HPDI Upper | ESS bulk | ESS tail | Rhat |
|---|---|---|---|---|---|---|---|---|
| 1 | $\log \lambda_0$ | -7.894 | 0.253 | -8.367 | -7.386 | 10341.749 | 8492.773 | 1.000 |
| 6 | $\log s'_{avf}$ | -7.824 | 0.608 | -9.318 | -6.805 | 11400.476 | 7020.896 | 1.000 |
| 7 | $\log s'_{dist}$ | -8.314 | 0.757 | -10.102 | -7.034 | 13407.794 | 8451.297 | 1.000 |
| 8 | $\log s'_{vor}$ | -8.096 | 0.795 | -9.884 | -6.772 | 12017.240 | 7417.397 | 1.001 |
| 9 | $\log s'_{fov}$ | -7.158 | 0.749 | -9.038 | -5.864 | 11416.267 | 6132.822 | 1.000 |
| 5 | $\lambda_0$ | 0.000 | 0.000 | 0.000 | 0.001 | 10341.736 | 8492.773 | 1.000 |
| 10 | $s_{avf}$ | 1.056 | 0.736 | 0.026 | 2.871 | 9874.471 | 7261.160 | 1.000 |
| 11 | $s_{dist}$ | 0.651 | 0.505 | 0.017 | 1.963 | 12071.244 | 8228.917 | 1.000 |
| 12 | $s_{vor}$ | 0.817 | 0.658 | 0.007 | 2.525 | 11335.848 | 7390.838 | 1.000 |
| 13 | $s_{fov}$ | 2.054 | 1.592 | 0.031 | 5.963 | 10890.753 | 5842.623 | 1.000 |
| 2 | $\beta_{\text{ILViavfmon}}$ | 1.694 | 0.550 | 0.626 | 2.739 | 12737.748 | 8886.308 | 1.000 |
| 3 | $\beta_{\text{ILVitimeHD}}$ | -1.812 | 0.407 | -2.658 | -1.075 | 16605.972 | 9118.880 | 1.000 |
| 4 | $\beta_{\text{ILVstimeHD}}$ | -1.777 | 0.641 | -3.048 | -0.511 | 16550.910 | 9659.682 | 1.000 |
| 14 | $\sigma_{ID,log(s'_{avf})}$ | 0.282 | 0.250 | 0.000 | 0.828 | 10212.305 | 6207.423 | 1.000 |
| 15 | $\sigma_{ID,log(s'_{dist})}$ | 0.302 | 0.266 | 0.000 | 0.877 | 9798.910 | 5451.645 | 1.000 |
| 16 | $\sigma_{ID,log(s'_{vor})}$ | 0.313 | 0.265 | 0.000 | 0.885 | 11149.311 | 6986.325 | 1.000 |
| 17 | $\sigma_{ID,log(s'_{fov})}$ | 0.296 | 0.260 | 0.000 | 0.874 | 10920.171 | 6098.853 | 1.000 |
| 18 | $\sigma_{ID,log(\lambda_0)}$ | 0.275 | 0.188 | 0.000 | 0.603 | 4301.880 | 4783.813 | 1.001 |
| 19 | $\%ST$ avf | 0.120 | 0.060 | 0.015 | 0.226 | 10924.553 | 7349.966 | 1.000 |
| 20 | $\%ST$ dist | 0.072 | 0.048 | 0.003 | 0.163 | 12523.974 | 8222.285 | 1.000 |
| 21 | $\%ST$ vor | 0.072 | 0.050 | 0.003 | 0.168 | 11579.004 | 7563.296 | 1.000 |
| 22 | $\%ST$ fov | 0.048 | 0.029 | 0.001 | 0.101 | 10901.990 | 6199.293 | 1.000 |

Table S8: Posterior summary statistics for model parameters from best fitting multi-network model (“distractor” model). Columns include parameter name, median value, median absolute deviation (MAD), the lower and upper bounds of the 95% highest posterior density estimates, the number of effective samples and Rhat score. Parameters are fit on the log scale, and thus we included $log(\lambda_0)$ and $log(s')$, as well as their standard interpretations. $\beta$ parameters are individual-level variables for the intrinsic rate (ILVi) and social rate (ILVs). $\sigma_{ID}$ parameters are the scale parameter for varying effects. $\%ST$ gives the estimated percent of events that occurred through social transmission.

| | Parameter | Median | MAD | HPDI Lower | HPDI Upper | ESS bulk | ESS tail | Rhat |
|---|---|---|---|---|---|---|---|---|
| 1 | $\log\lambda_0$ | -7.774 | 0.243 | -8.263 | -7.304 | 7394.853 | 8436.078 | 1.001 |
| 2 | $\log s'_{avf}$ | -7.058 | 0.320 | -7.792 | -6.469 | 9240.513 | 5900.764 | 1.000 |
| 8 | $\lambda_0$ | 0.000 | 0.000 | 0.000 | 0.001 | 7394.858 | 8436.078 | 1.001 |
| 7 | $s_{avf}$ | 2.025 | 0.911 | 0.548 | 4.344 | 7000.507 | 6729.403 | 1.000 |
| 3 | $\beta_{\text{ILViavfmon}}$ | 1.668 | 0.517 | 0.639 | 2.654 | 9898.232 | 8620.630 | 1.000 |
| 4 | $\beta_{\text{ILVitimeHD}}$ | -1.946 | 0.389 | -2.745 | -1.187 | 11973.234 | 9116.301 | 1.000 |
| 5 | $\beta_{\text{ILVstimeHD}}$ | -1.454 | 0.665 | -2.824 | -0.136 | 10583.021 | 8897.872 | 1.001 |
| 9 | $\sigma_{ID,log(s'_{avf})}$ | 0.227 | 0.199 | 0.000 | 0.673 | 7487.552 | 6035.554 | 1.000 |
| 10 | $\sigma_{ID,log(\lambda_0)}$ | 0.346 | 0.183 | 0.001 | 0.656 | 4101.825 | 5274.869 | 1.001 |
| 11 | $\%ST$ avf | 0.266 | 0.050 | 0.164 | 0.359 | 9985.390 | 6564.829 | 1.000 |

Table S9: Posterior summary statistics for model parameters from best fitting single network ("distractor" model). Columns include parameter name, median value, median absolute deviation (MAD), the lower and upper bounds of the 95% highest posterior density estimates, the number of effective samples and Rhat score. Parameters are fit on the log scale, and thus we included $log(\lambda_0)$ and $log(s')$, as well as their standard interpretations. $\beta$ parameters are individual-level variables for the intrinsic rate (ILVi) and social rate (ILVs). $\sigma_{ID}$ parameters are the scale parameter for varying effects.

| Behaviour | Description |
|---|---|
| Duration foveating | Time spent foveating on the monitor object (either the target or the distractor) during the time interval. |
| Duration head-up | Time spent in a head-up position and not distracted by any other activity (feeding, courting or grooming) during the time interval. |
| Became head-up | Number of times the individual switches from a feeding state to a head-up state (not feeding, courting nor grooming), over the time interval. |
| Frequency of saccades | Number of head-scan saccades (fast head movements) made by the individual over the time interval. |
| Duration walking | Time spent walking (locomotion speed above 0.2 m/s) during the time interval. |
| Duration running away from the monitor | Time spent walking fast (locomotion speed above 0.6 m/s), and away from the monitor displaying the looming stimulus (target), during the time interval. |
| Duration flying | Time spent flying (locomotion speed above 2 m/s) during the time interval. |

Table S10: Description of the different behavioural cues used as alternatives for the foveation on the monitor.

| | Parameter | Median | MAD | HPDI Lower | HPDI Upper | ESS bulk | ESS tail | Rhat |
|---|---|---|---|---|---|---|---|---|
| 1 | $\log\lambda_0$ | -7.465 | 0.260 | -7.990 | -6.955 | 13421.814 | 9020.536 | 1.000 |
| 7 | $\log s'_{avf}$ | -6.960 | 0.434 | -7.905 | -6.118 | 6988.943 | 7580.411 | 1.000 |
| 8 | $\log s'_{dist}$ | -9.327 | 0.820 | -11.160 | -7.919 | 13843.989 | 8764.622 | 1.001 |
| 9 | $\log s'_{vor}$ | -8.720 | 0.833 | -10.635 | -7.255 | 11043.550 | 8876.935 | 1.001 |
| 10 | $\log s'_{fov}$ | -8.062 | 0.937 | -10.111 | -6.514 | 16236.453 | 8830.124 | 1.001 |
| 6 | $\lambda_0$ | 0.001 | 0.000 | 0.000 | 0.001 | 13421.828 | 9020.536 | 1.000 |
| 11 | $s_{avf}$ | 1.667 | 0.811 | 0.289 | 3.540 | 7150.241 | 7787.977 | 1.000 |
| 12 | $s_{dist}$ | 0.154 | 0.126 | 0.004 | 0.523 | 13647.645 | 9082.425 | 1.001 |
| 13 | $s_{vor}$ | 0.285 | 0.237 | 0.005 | 0.981 | 11259.605 | 9105.560 | 1.001 |
| 14 | $s_{fov}$ | 0.549 | 0.486 | 0.007 | 2.088 | 15669.148 | 9294.116 | 1.000 |
| 2 | $\beta_{\text{ILViavfmon}}$ | 3.044 | 0.495 | 2.060 | 3.984 | 19009.121 | 10506.618 | 1.000 |
| 3 | $\beta_{\text{ILVitimeHD}}$ | -2.100 | 0.352 | -2.790 | -1.423 | 15664.504 | 10459.455 | 1.000 |
| 4 | $\beta_{\text{ILVstimeHD}}$ | -2.553 | 0.618 | -3.791 | -1.340 | 14201.477 | 8499.354 | 1.001 |
| 5 | $\beta_{\text{ILVmtrial}}$ | 0.021 | 0.023 | -0.024 | 0.064 | 16852.324 | 9713.919 | 1.000 |
| 20 | $\%ST$ avf | 0.231 | 0.069 | 0.098 | 0.381 | 6138.161 | 6724.869 | 1.000 |
| 21 | $\%ST$ dist | 0.024 | 0.018 | 0.001 | 0.066 | 12945.015 | 9307.932 | 1.001 |
| 22 | $\%ST$ vor | 0.039 | 0.029 | 0.001 | 0.110 | 9761.279 | 8792.997 | 1.000 |
| 23 | $\%ST$ fov | 0.014 | 0.012 | 0.000 | 0.044 | 15622.597 | 8710.661 | 1.000 |
| 15 | $\sigma_{ID,log(s'_{avf})}$ | 0.738 | 0.302 | 0.044 | 1.280 | 3779.241 | 4136.096 | 1.001 |
| 16 | $\sigma_{ID,log(s'_{dist})}$ | 0.338 | 0.292 | 0.000 | 0.965 | 10911.916 | 6449.890 | 1.000 |
| 17 | $\sigma_{ID,log(s'_{vor})}$ | 0.370 | 0.315 | 0.000 | 1.012 | 9895.114 | 6975.040 | 1.000 |
| 18 | $\sigma_{ID,log(s'_{fov})}$ | 0.320 | 0.281 | 0.000 | 0.932 | 13179.931 | 6621.592 | 1.000 |
| 19 | $\sigma_{ID,log(\lambda_0)}$ | 0.382 | 0.173 | 0.001 | 0.686 | 4392.144 | 4355.932 | 1.000 |

Table S11: Posterior summary statistics for model parameters from multi-network model, refit with trial as a multiplicative effect. Trial was not estimated to affect either asocial or social learning pathways. Columns include parameter name, median value, median absolute deviation (MAD), the lower and upper bounds of the 95% highest posterior density estimates, the number of effective samples and Rhat score. Parameters are fit on the log scale, and thus we included $log(\lambda_0)$ and $log(s')$, as well as their standard interpretations. $\beta$ parameters are individual-level variables for the intrinsic rate (ILVi) and social rate (ILVs). $\sigma_{ID}$ parameters are the scale parameter for varying effects.

| | Parameter | Median | MAD | HPDI Lower | HPDI Upper | ESS bulk | ESS tail | Rhat |
|---|---|---|---|---|---|---|---|---|
| 1 | $\log\lambda_0$ | -7.348 | 0.230 | -7.827 | -6.894 | 10570.151 | 8870.138 | 1.000 |
| 6 | $\log s'_{avf}$ | -6.753 | 0.365 | -7.677 | -6.092 | 6352.056 | 5789.011 | 1.000 |
| 7 | $\log s'_{dist}$ | -9.393 | 0.858 | -11.289 | -7.853 | 12142.638 | 8272.526 | 1.001 |
| 8 | $\log s'_{vor}$ | -8.726 | 0.926 | -10.824 | -7.221 | 9221.939 | 7965.088 | 1.001 |
| 9 | $\log s'_{fov}$ | -8.166 | 0.993 | -10.309 | -6.458 | 13734.166 | 8280.800 | 1.000 |
| 5 | $\lambda_0$ | 0.001 | 0.000 | 0.000 | 0.001 | 10570.148 | 8870.138 | 1.000 |
| 10 | $s_{avf}$ | 1.801 | 0.842 | 0.394 | 3.861 | 6488.427 | 6038.167 | 1.000 |
| 11 | $s_{dist}$ | 0.129 | 0.109 | 0.002 | 0.469 | 11932.903 | 7986.829 | 1.000 |
| 12 | $s_{vor}$ | 0.252 | 0.229 | 0.002 | 0.943 | 9622.864 | 8112.329 | 1.000 |
| 13 | $s_{fov}$ | 0.440 | 0.408 | 0.005 | 1.773 | 13656.080 | 8449.601 | 1.000 |
| 2 | $\beta_{\text{ILViavfmon}}$ | 3.110 | 0.498 | 2.166 | 4.084 | 13444.944 | 9906.992 | 1.000 |
| 3 | $\beta_{\text{ILVitimeHD}}$ | -2.151 | 0.352 | -2.884 | -1.489 | 15388.353 | 9277.436 | 1.000 |
| 4 | $\beta_{\text{ILVstimeHD}}$ | -2.493 | 0.598 | -3.775 | -1.395 | 14669.324 | 9076.093 | 1.001 |
| 19 | $\%ST$ avf | 0.286 | 0.059 | 0.153 | 0.393 | 5265.552 | 4967.287 | 1.000 |
| 20 | $\%ST$ dist | 0.023 | 0.018 | 0.000 | 0.069 | 11374.025 | 8381.994 | 1.001 |
| 21 | $\%ST$ vor | 0.040 | 0.033 | 0.001 | 0.119 | 7869.105 | 7471.220 | 1.001 |
| 22 | $\%ST$ fov | 0.013 | 0.012 | 0.000 | 0.044 | 13424.025 | 8207.994 | 1.000 |
| 14 | $\sigma_{ID,log(s'_{avf})}$ | 0.751 | 0.295 | 0.042 | 1.259 | 3511.959 | 3037.592 | 1.001 |
| 15 | $\sigma_{ID,log(s'_{dist})}$ | 0.337 | 0.289 | 0.000 | 0.960 | 10538.134 | 6164.586 | 1.000 |
| 16 | $\sigma_{ID,log(s'_{vor})}$ | 0.363 | 0.317 | 0.000 | 1.018 | 9631.233 | 7315.198 | 1.000 |
| 17 | $\sigma_{ID,log(s'_{fov})}$ | 0.319 | 0.274 | 0.000 | 0.915 | 11126.870 | 6466.762 | 1.000 |
| 18 | $\sigma_{ID,log(\lambda_0)}$ | 0.368 | 0.175 | 0.000 | 0.670 | 3710.893 | 3419.833 | 1.000 |

Table S12: Posterior summary statistics for model parameters from best-fitting multi-network model, refit with a prior more skeptical of social transmission ($log(\lambda_0) \sim (-5,2)$, $log(s') \sim (-5,2)$, thus $s \approx 0.36$). Estimates are similar to priors that assume $s = 1$. Estimates are similar, but with additional shrinkage. Columns include parameter name, median value, median absolute deviation (MAD), the lower and upper bounds of the 95% highest posterior density estimates, the number of effective samples and Rhat score. Parameters are fit on the log scale, and thus we included $log(\lambda_0)$ and $log(s')$, as well as their standard interpretations. $\beta$ parameters are individual-level variables for the intrinsic rate (ILVi) and social rate (ILVs). $\sigma_{ID}$ parameters are the scale parameter for varying effects.
